# The evolutionary plasticity of chromosome metabolism allows adaptation to DNA replication stress

**DOI:** 10.1101/770859

**Authors:** Marco Fumasoni, Andrew W. Murray

## Abstract

Chromosome metabolism is defined by the pathways that collectively maintain the genome, including chromosome replication, repair and segregation. Because aspects of these pathways are conserved, chromosome metabolism is considered resistant to evolutionary change. We used the budding yeast, *Saccharomyces cerevisiae*, to investigate the evolutionary plasticity of chromosome metabolism. We experimentally evolved cells constitutively experiencing DNA replication stress caused by the absence of Ctf4, a protein that coordinates the activities at replication forks. Parallel populations adapted to replication stress, over 1000 generations, by acquiring multiple, successive mutations. Whole-genome sequencing and testing candidate mutations revealed adaptive changes in three aspects of chromosome metabolism: DNA replication, DNA damage checkpoint and sister chromatid cohesion. Although no gene was mutated in every population, the same pathways were sequentially altered, defining a functionally reproducible evolutionary trajectory. We propose that this evolutionary plasticity of chromosome metabolism has important implications for genome evolution in natural populations and cancer.

## Introduction

The central features of many fundamental biological processes, such as the mechanism of DNA, RNA and protein synthesis, have been conserved since the last common ancestor of all extant organisms. Many of the proteins involved in these processes are essential, and the complex molecular interactions between them have been argued to constrain the evolution of both the processes and the proteins that carry them out (Wilson, Carlson, and White 1977; Fraser et al. 2002).

DNA replication is one of the most conserved cellular processes. Replication requires multiple enzymes that catalyze individual reactions such as unwinding the double helix, priming replication, and synthesizing new DNA strands (O‘Donnell, Langston, and Stillman 2013). A common feature of replication is the organization of these enzymatic activities in multi-molecular complexes called replisomes, whose function is to coordinate the simultaneous synthesis of DNA from the two anti-parallel template strands (Yao and O‘Donnell 2016). Replisomes need to be tightly regulated to integrate replication with other essential aspects of chromosome metabolism such as DNA repair and chromosome segregation (Branzei and Foiani 2010; Bell and Labib 2016). This regulation is critical in eukaryotes, where the presence of multiple replication origins requires the coordination of several replisomes simultaneously travelling along the same DNA molecule (Dewar and Walter 2017; Siddiqui, On, and Diffley 2013).

The temporal and physical interactions between the enzymatic machinery that performs the different steps of DNA replication are remarkably conserved. Nevertheless, differences in many features of DNA replication have been reported: the number of replisome subunits is higher in eukaryotes than in bacteria, possibly to account for the higher complexity of eukaryotic genomes (McGeoch and Bell 2008). Some subunits are only found in some eukaryotic species (Y. Liu, Richards, and Aves 2009; Aves, Liu, and Richards 2012). Notably, there are also biochemical variations in important features, such as the helicase, which encircles the leading strand in eukaryotes and the lagging strand in prokaryotes (McGeoch and Bell 2008), or differences in the regulation of DNA replication by the machinery that drives the cell cycle progression (Cross, Buchler, and Skotheim 2011; Siddiqui, On, and Diffley 2013; Parker, Botchan, and Berger 2017).

These differences reveal that although the DNA replication module performs biochemically conserved reactions, its features can change during evolution. This observation poses an apparent paradox: how can such an important process change during evolution without killing cells? One hypothesis is that because so many replication proteins are essential, the observed differences can only be obtained by extremely slow evolutionary processes that require many successive mutations of small effect and happen over millions of generations. Alternatively, the DNA replication module could accommodate substantial changes within hundreds or thousands of generations, but such events would have to be rare to explain the overall conservation of DNA replication.

To distinguish between these two hypotheses, we followed the evolutionary response to a genetic perturbation of DNA replication. Characterizing evolutionary responses to genetic perturbations has informed studies of functional modules (Rojas Echenique et al. 2019; Filteau et al. 2015; Harcombe, Springman, and Bull 2009), challenged the notion that particular genes are essential (Rancati et al. 2018; G. Liu et al. 2015), and revealed that initial genotypes can determine evolutionary trajectories (Szamecz et al. 2014; Rojas Echenique et al. 2019; Lind, Farr, and Rainey 2015).

We followed the evolutionary response of *S. cerevisiae* to DNA replication stress, an overall perturbation of DNA replication that interferes with chromosome metabolism, reduces cell viability, and induces genetic instability (Zeman and Cimprich 2014; Muñoz and Méndez 2016). DNA replication stress has been implicated in both cancer progression and aging (Burhans and Weinberger 2007; Gaillard, García-Muse, and Aguilera 2015) but despite studies investigating the direct effect of replication stress on cell physiology, its evolutionary consequences are unknown. We induced replication stress by removing an important but non-essential component of the DNA replication module, Ctf4, which coordinates activities at the replisome (Villa et al. 2016). We then evolved eight *ctf4 Δ*populations for 1000 generations, exploiting the ability of experimental evolution (Barrick and Lenski 2013) to identify, analyze, and compare the mutations that create parallel evolutionary trajectories to increase fitness (Laan, Koschwanez, and Murray 2015; Koschwanez, Foster, and Murray 2013; Wildenberg and Murray 2014).

We found that populations recover from the fitness defect induced by DNA replication stress. Genetic analysis revealed that their adaptation is driven by mutations that change conserved features in three modules involved in chromosome metabolism: DNA replication, the DNA damage checkpoint, and sister chromatid cohesion. These mutations arise sequentially and collectively allow cells to approach the fitness of their wild-type ancestors within 1000 generations of evolution. The molecular basis of these adaptive strategies and their epistatic interactions produce a mechanistic model of the evolutionary adaptation to replication stress. Our results reveal the short-term evolutionary plasticity of chromosome metabolism. We discuss the consequences of this plasticity for the evolution of species in the wild and cancer progression.

## Results

### Adaptation to DNA replication stress is driven by mutations in chromosome metabolism

Replication stress refers to the combination of the defects in DNA metabolism and the cellular response to these defects in cells whose replication has been substantially perturbed (Macheret and Halazonetis 2015). Problems in replication can arise at the sites of naturally occurring or experimentally induced lesions and can cause genetic instability (Muñoz and Méndez 2016). We asked how cells evolve to adapt to constitutive DNA replication stress.

Previous work has induced replication stress by using chemical treatments or genetic perturbations affecting factors involved in DNA replication (Zheng et al. 2016; Tkach et al. 2012; Mazouzi et al. 2016). To avoid evolving resistance to drugs or the reversion of point mutations that induce replication stress, we chose instead to remove *CTF4*, a gene encoding an important, but non-essential, component of the DNA replication machinery. Ctf4 is a homo-trimer, that serves as a structural hub within the replisome and coordinates different aspects of DNA replication by binding the replicative helicase, the primase, and other factors recruited to the replication fork (Figure 1A, Gambus et al., 2009; Samora et al., 2016; Simon et al., 2014; Tanaka et al., 2009; Villa et al., 2016). In the absence of Ctf4, cells experience several problems in fork progression leading to the accumulation of defects commonly associated with DNA replication stress (Muñoz and Méndez 2016), such as single-stranded DNA gaps and altered replication forks (Fumasoni et al. 2015; Abe et al. 2018; Kouprina et al. 1992).

**Figure 1.**
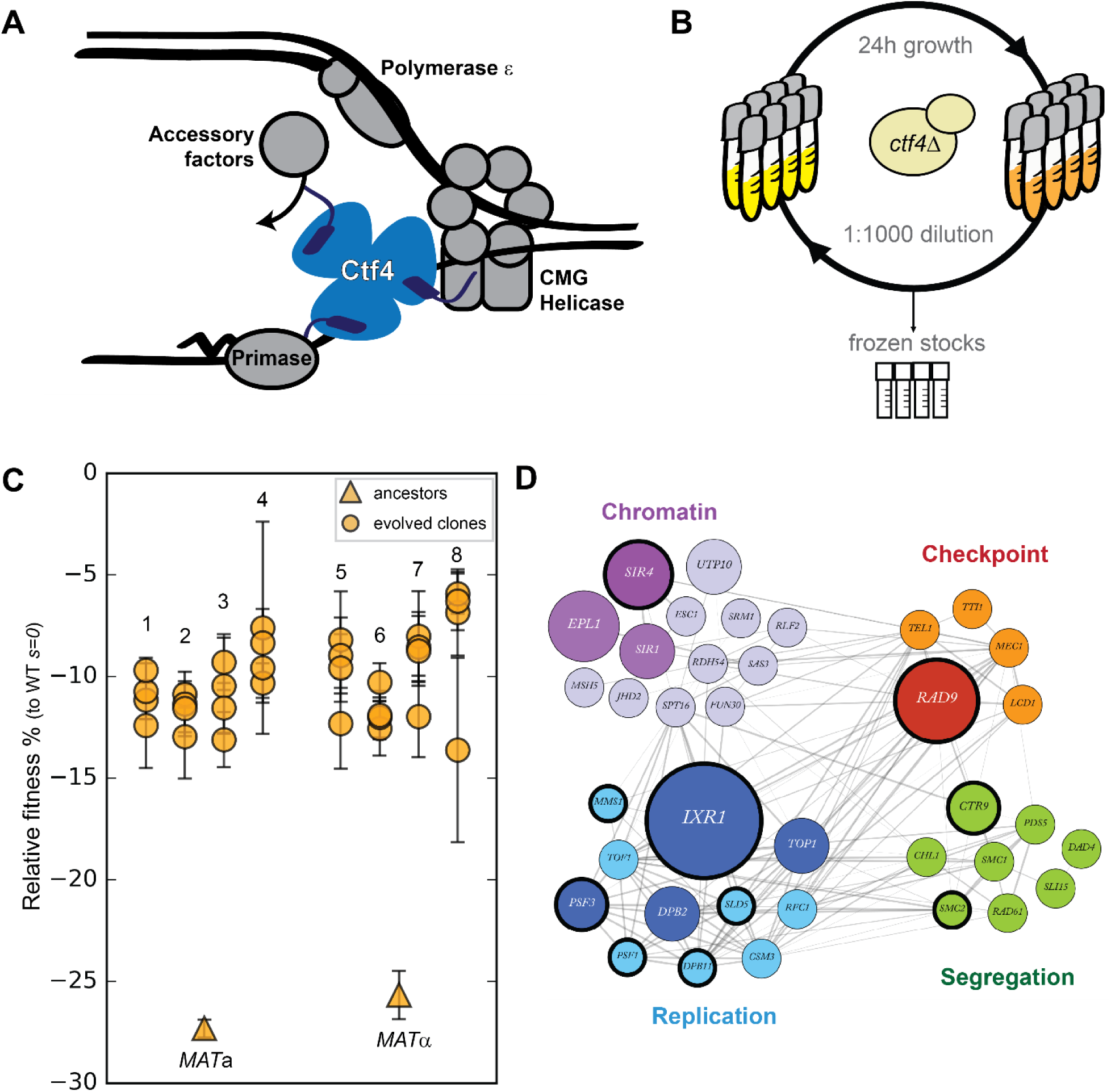
Fast evolutionary adaptation to DNA replication stress. **(A)** Schematic representation of the replisome focused on the role of Ctf4 in coordinating the replicative helicase, primase, and other factors. **(B)** The experimental evolution scheme: independent colonies of *ctf4Δ S. cerevisiae* were inoculated in rich media, grown to saturation, and diluted 1:1000 in fresh media for a total of 100 cycles (1000 generations). Populations samples were saved every 50 generations for future analysis. **(C)** Fitness of the *ctf4Δ* ancestor strains and of 32 evolved clones isolated from the 8 (labeled 1 through 8) populations derived from them, relative to wt cells (*s*=0). Error bars represent standard deviations. *MAT****a*** and *MATα* refer to the strain sex. **(D)** Simplified representation of the modules enriched in putative adaptive mutations, found in evolved clones. Grey lines represent evidence of genetic and physical interactions from the literature (https://string-db.org). Node diameter is proportional to the number of populations in which the gene was mutated. Selection on darker nodes was statistically significant. Nodes surrounded with a bold circle are genes in which mutations were found to strongly correlate with the evolved phenotype by bulk segregant analysis.

We generated *ctf4*Δ and wild type (WT) ancestor strains by sporulating a heterozygous *CTF4/ctf4*Δ diploid. As previously reported (Miles and Formosa 1992; Kouprina et al. 1992), *ctf4*Δ cells display severe growth defects, which we quantified as a fitness decrease of approximately 25% relative to WT (Figure 1C). We then evolved eight parallel populations of each genotype for 1000 generations by serial dilutions in rich media, freezing population samples every 50 generations (Figure 1B). Under this regime, spontaneous mutations that increase cellular fitness and survive genetic drift will be selected and spread asexually within the populations (Jerison and Desai 2015; Venkataram et al. 2016; Levy et al. 2015). At the end of the experiment, we asked whether cells had recovered from the fitness decrease induced by replication stress by measuring the fitness of the evolved *ctf4*Δ and WT populations. Expressing the results as a percentage of the fitness of the WT ancestor, the evolved WT populations increased their fitness by an average of 4.0±0.3% (Figure S1A), a level similar to previous experiments (Lang et al. 2013; Buskirk, Peace, and Lang 2017). In contrast, we found that the fitness of the evolved *ctf4Δ* populations rose by 17±0.2% (Figure S1A). Clones isolated from these populations showed similar fitness increases (Figure 1C).

To understand this evolutionarily rapid adaptation to constitutive replication stress, we whole-genome sequenced all the final evolved populations as well as 32 individual clones (4 from each of the evolved populations) isolated from the *ctf4Δ* lineages. During experimental evolution, asexual populations accumulate two types of mutations: adaptive mutations that increase their fitness and neutral or possibly mildly deleterious mutations that hitchhike with the adaptive mutations (Table S1). To distinguish between these mutations, we used a combination of statistical and experimental approaches. First, we inferred that mutations in a gene were adaptive if the gene was mutated more frequently than expected by chance across our parallel and independent populations (Table S2). Second, we performed bulk segregant analysis on selected evolved clones. This technique takes advantage of sexual reproduction, followed by selection, to separate causal and hitchhiking mutations. In this case, mutations that segregate strongly with the evolved phenotype are assumed to be adaptive (Figure S1B). We combined these two lists of mutated genes and looked for enriched gene ontology (GO) terms. This analysis revealed an enrichment of genes implicated in several aspect of chromosome metabolism (Table S3). Among the genes associated with these terms, many are involved in four functional modules: DNA replication, chromosome segregation (including genes involved in sister chromatid linkage and spindle function), cell cycle checkpoint and chromatin remodeling (Figure S1C). The genes in these modules that were mutated in the evolved clones are shown, grouped by function, in Figure 1D.

### Loss of the DNA damage checkpoint shortens G2/M

We found several mutations affecting genes involved in cell-cycle checkpoints (Figure 2B). Checkpoints are feedback control mechanisms that induce cell-cycle delays in response to defects that reflect the failure to complete important process and thus guarantee the proper sequence of events required for cell division (Elledge 1996; Murray 1992). Three such delays have been characterized. In yeast, the first prevent cells from entering S-phase in response to DNA damage occurring in G1. A second slow progress through S-phase in response to problems encountered during DNA synthesis. The third can instead delay sister chromatid separation (anaphase) and the exit from mitosis in response to DNA damage incurred after cells enter S-phase or defects in chromosome attachment to the mitotic spindle (Figure 2A, Murray, 1994).

**Figure 2.**
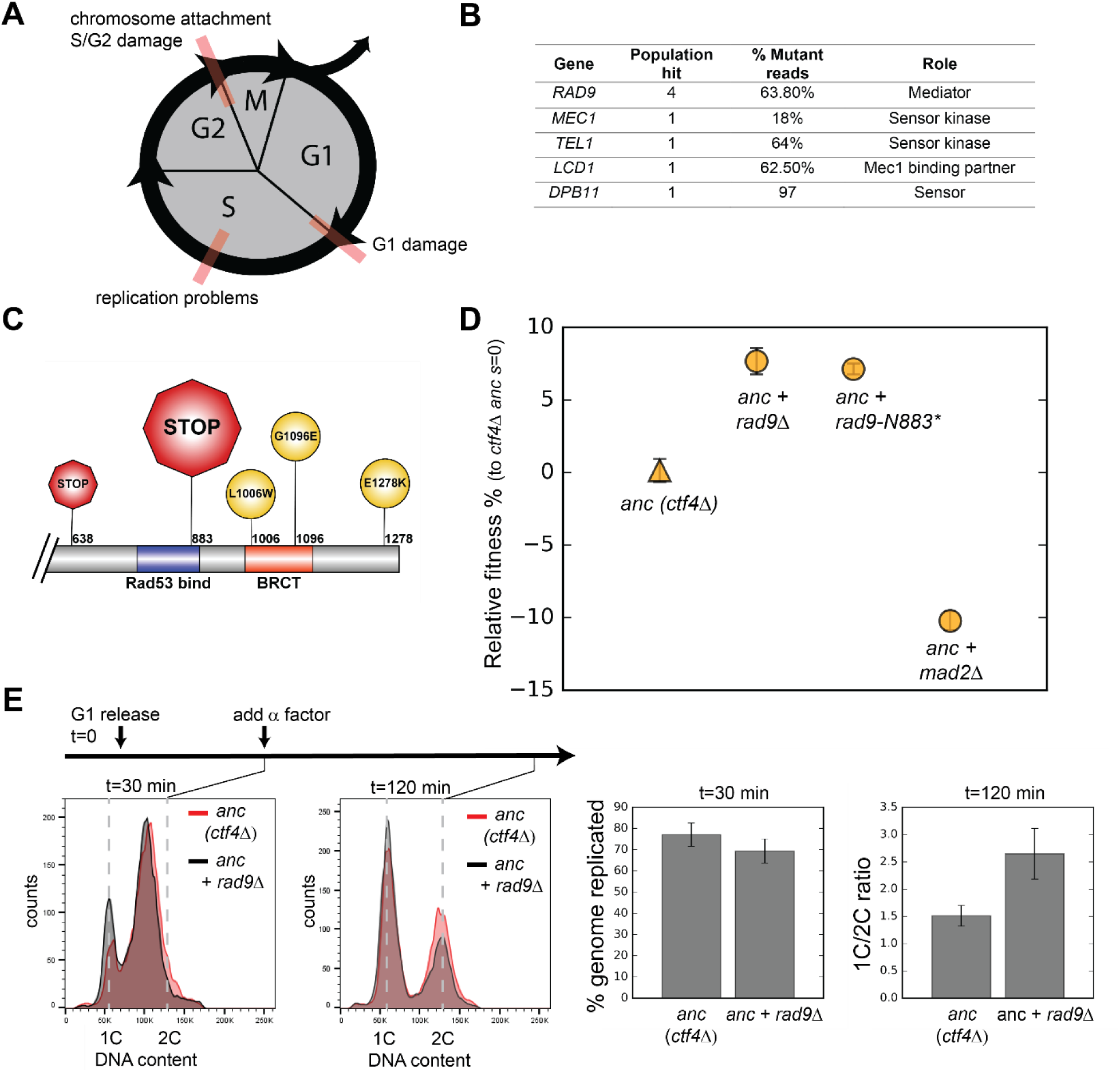
Checkpoint mutations cause a faster G2/M transition in evolved cells. **(A)** Schematic representation of cell cycle progression. The transitions delayed by various checkpoints are highlighted in red. **(B)** List of checkpoint genes mutated in evolved clones and their role in the signaling cascade. ‘Populations hit’ refers to the number of populations where the gene was mutated. ‘% Mutant reads’ was calculated as the average of the mutant read frequencies in the different populations where the mutation was detected. **(C)** Schematic of the C-terminal region of Rad9 that was affected by mutations in evolved clones. The diameter of the symbol is proportional to the number of populations where the mutation was detected. Note that both stop codons resulted from an upstream frameshift. Two populations contained more than one distinct *RAD9* mutations. **(D)** The fitness of *ctf4Δ* strains carrying two reconstructed mutations in the DNA damage checkpoint (*rad9Δ* and *rad9K883\**) and an engineered inactivation of the spindle checkpoint (*mad2Δ*) relative to the *ctf4Δ* ancestors (*ctf4Δ anc, s*=0). Error bars represent standard deviations. **(E)** Cell cycle profiles of *ctf4Δ* ancestor and *ctf4Δ rad9Δ* cells at two time points during a synchronous cell cycle. Cell were arrested in G1 and subsequently released synchronously into S-phase. Time points taken at 30 min and 120 min after the release are shown. *α*-factor was added 30 min after release to prevent cells entering a second cell cycle and thus ensure that 2C cells at the 120 min measurement resulted from a G2 delay rather than progress through a second cell cycle. The percentage of genome replicated at 30 min was calculated based on the cell cycle profile.1C/2C ratios were calculated based on the height of the respective 1C and 2C peaks at 120 min.

The genes listed in Figure 2B are implicated at different levels in either the replication or mitotic delays (Figure S2B, Pasero and Vindigni, 2017). The most frequently mutated gene, *RAD9*, encodes an important component of the DNA damage checkpoint, which is required to slow DNA synthesis and delay anaphase in response to DNA lesions (Weinert and Hartwell 1988). Four out of the five mutations in *RAD9* produced early stop codons, or radical amino acid substitutions in the BRCT domain, which is essential for Rad9‘s function (Figure 2C, S2A, Soulier and Lowndes, 1999), arguing that inactivation of Rad9 was repeatedly and independently selected for during evolution. To test this hypothesis, we engineered the most frequently occurring mutation (*2628* +*A*, a frameshift mutation leading to a premature stop codon K883*) into the ancestral *ctf4Δ* strain (*ctf4Δ* anc). We suspect that the high frequency of this mutation is due to the presence of a run of 11 As, a sequence that is known to be susceptible to loss or gain of a base during DNA replication. This mutation (Figure 2C, S2A) produced a fitness increase very similar to the one caused by deleting the entire gene (Figure 2D). We conclude that inactivation of Rad9 is adaptive in the absence of Ctf4.

We asked if the removal of Rad9 eliminated a cell cycle delay caused by the absence of Ctf4. In the *ctf4*Δ ancestor, *rad9*Δ does, indeed, decrease the fraction of cells with a 2C DNA content (the DNA content in G2 and mitosis) observed in asynchronously growing *ctf4*Δ cells (Tanaka et al., 2009). This observation suggests that the interval between the end of DNA replication and cell division decreases in *ctf4*Δ *rad9*Δ cells. The spindle checkpoint, which blocks anaphase in response to defects in mitotic spindle assembly, can also delay chromosome segregation in cells (Li and Murray 1991). But although deleting *MAD2*, a key spindle checkpoint component, also decreases the interval between replication and division in *ctf4*Δ cells (Hanna et al. 2001), it reduces rather than increases the fitness of the *ctf4*Δ ancestor (Figure 2D). These results suggest that ignoring some defects in *ctf4*Δ cells, such as those that activate the DNA damage checkpoint, improves fitness, whereas ignoring others, such as defects in chromosome alignment on the spindle, reduces fitness.

Problems encountered during DNA synthesis also activate the replication checkpoint, which inhibits DNA replication to prevent further lesions (Zegerman and Diffley 2009, 2010). As many proteins involved in the DNA damage checkpoint are shared with the replication checkpoint (Figure S2B, Pardo et al., 2017), we followed a single synchronous cell-cycle to ask whether the fitness benefits conferred by *RAD9* deletion were due to a faster progression through S-phase or faster progress through mitosis. Loss of Rad9 in *ctf4Δ* cells did not accelerate S-phase, but it did lead to faster passage through mitosis as revealed by a reduced fraction of 2C cells (Figure 2E).

To separate the role of the replication and DNA damage checkpoints, we genetically manipulated targets of the checkpoints whose phosphorylation delays either anaphase (Pds1, Wang et al., 2001) or the completion of replication (Sld3 and Dbf4, Zegerman & Diffley, 2010, Figure S2B). Fitness measurement in these mutants (*pds1-m9* or the double mutant *sld3-A/dbf4-4A*) showed that while decreasing the mitotic delay in ancestral *ctf4Δ* cells was beneficial, a faster S-phase was highly detrimental (Figure S2C). Collectively, these results show that the specific absence of a DNA damage-induced delay of anaphase, rather than generic cell-cycle acceleration, is adaptive in *ctf4Δ* cells experiencing replication stress.

### Amplification of cohesin loader genes improves sister chromatid cohesion

We examined the evolved clones for changes in the copy number across the genome (DNA copy number variations, CNVs). Several clones showed segmental amplifications, defined as an increase in the copy number of a defined chromosomal segment (Figure S3A). The most common (17 out of 32 sequenced clones) was the amplification of a 50-100 kb region of chromosome IV (chrIV). In addition to this segmental amplification on chromosome IV, evolved clone EVO2-10 also carried an extra copy of a portion of chromosome V (chrV, Figure 3A). Amongst the genes affected by these two CNVs are *SCC2* and *SCC4* on the amplified portions of chromosomes IV and V respectively. These two genes encode the two subunits of the cohesin loader complex, which loads cohesin rings on chromosomes to ensure sister chromatid cohesion until anaphase (Figure S3C, Ciosk et al., 2000; Michaelis et al., 1997). The amplification of *SCC2* and *SCC4*, together with the other genes altered by point mutations in our evolved clones (Figure 3B), strongly suggest that the absence of Ctf4 selects for mutations that affect the linkage between sister chromatids.

**Figure 3.**
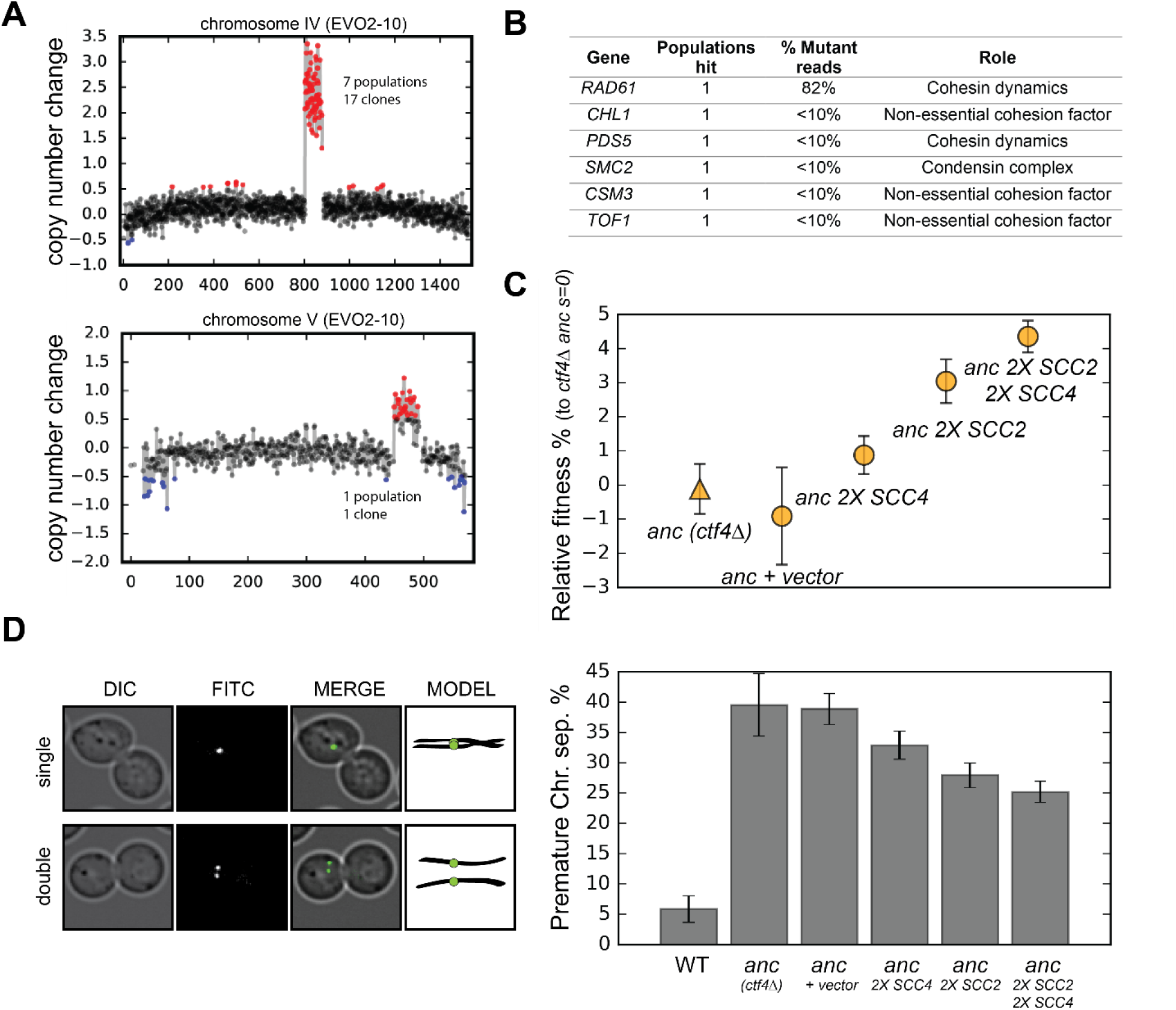
Amplification of cohesin loader genes. **(A)** Copy number variations (CNVs) affecting chromosome IV and chromosome V in clone EVO2-10. Copy number change refers to the fragment‘s gain or loss during the evolution experiment (i.e. +1 means that one copy was gained). Red highlights gains, blue highlights losses. **(B)** List of genes involved in chromosome segregation that were mutated in evolved clones, and their respective role in the process. ‘populations hit’ is the number of populations where the gene was found mutated. ‘% Mutant reads’ was calculated as the average of the mutant read frequencies in the different populations where the mutation was detected. **(C)** Fitness of ancestral, *ctf4Δ* strains that carry chromosomally integrated extra copies of cohesin loader genes, relative to the *ctf4Δ* ancestor (*s*=0). Error bars represent standard deviations. **(D)** Premature chromatid separation assay: Cells which contained a chromosome marked by a GFP dot (Lac repressor-GFP binding to an array of LacO sites) were arrested in metaphase and visualized under the microscope. The number of dots reports on premature sister chromatid separation. Two sister chromatids that are still linked to each other produce a single fluorescent dot (single, left panel), while cells whose sister chromatids have separated contain two distinguishable dots (double, left panel). Quantitation of premature sister chromatid separation in cells carrying extra copies of cohesin loader genes (right panel).

*CTF4* was originally identified because mutants in this gene reduced the fidelity of chromosome transmission (CTF = chromosome transmission fidelity, Spencer et al., 1990); later studies showed that this defect was due to premature sister chromatid separation, which resulted in increased chromosome loss at cell division (Hanna et al. 2001).

We hypothesized that the segmental amplifications of chrIV and chrV were selected to increase the amount of the cohesin loading complex. To test this idea, we reintroduced a second copy of these genes in a *ctf4*Δ ancestor. As predicted by the more frequent amplification *SCC2*, we found that an extra copy of *SCC4* increased fitness by less than 2%, whereas an extra copy of *SCC2*, or an extra copy of both *SCC2* and *SCC4* increased fitness by 4-5% (Figure 3C). We examined cells arrested in mitosis to measure the extent of premature sister chromatid separation in the same strains. Adding extra copies of the cohesin loader subunits improved sister chromatid cohesion (Figure 3D) and the amplitude of the improvement in sister cohesion for different strains had the same rank order as their increase in fitness (Figure 3C). We conclude that the increased copy number of the cohesin loader subunits is adaptive and alleviates the cohesion defects induced by the lack of Ctf4.

### Altered replication dynamics promote DNA synthesis in late replication zones

We found mutations in several genes involved in DNA replication (Figure 4A). Among these, we found four independent mutations (Figure 4B) that altered three different subunits of the replicative CMG (Cdc45, MCM, GINS) helicase (Moyer, Lewis, and Botchan 2006; Labib and Gambus 2007). The CMG helicase is bound in vivo by Ctf4 through the GINS subunit Sld5 (Simon et al. 2014). This binding allows Ctf4 to coordinate the helicase‘s progression with primase, which synthesizes the primers for lagging strand DNA synthesis, and other factors recruited behind the replication fork (Figure 1A, Samora et al., 2016; Villa et al., 2016). A CMG helicase mutation found in one of the evolved clones, *sld5-E130K*, increased the fitness of the ancestral *ctf4*Δ strain (Figure 4C).

**Figure 4.**
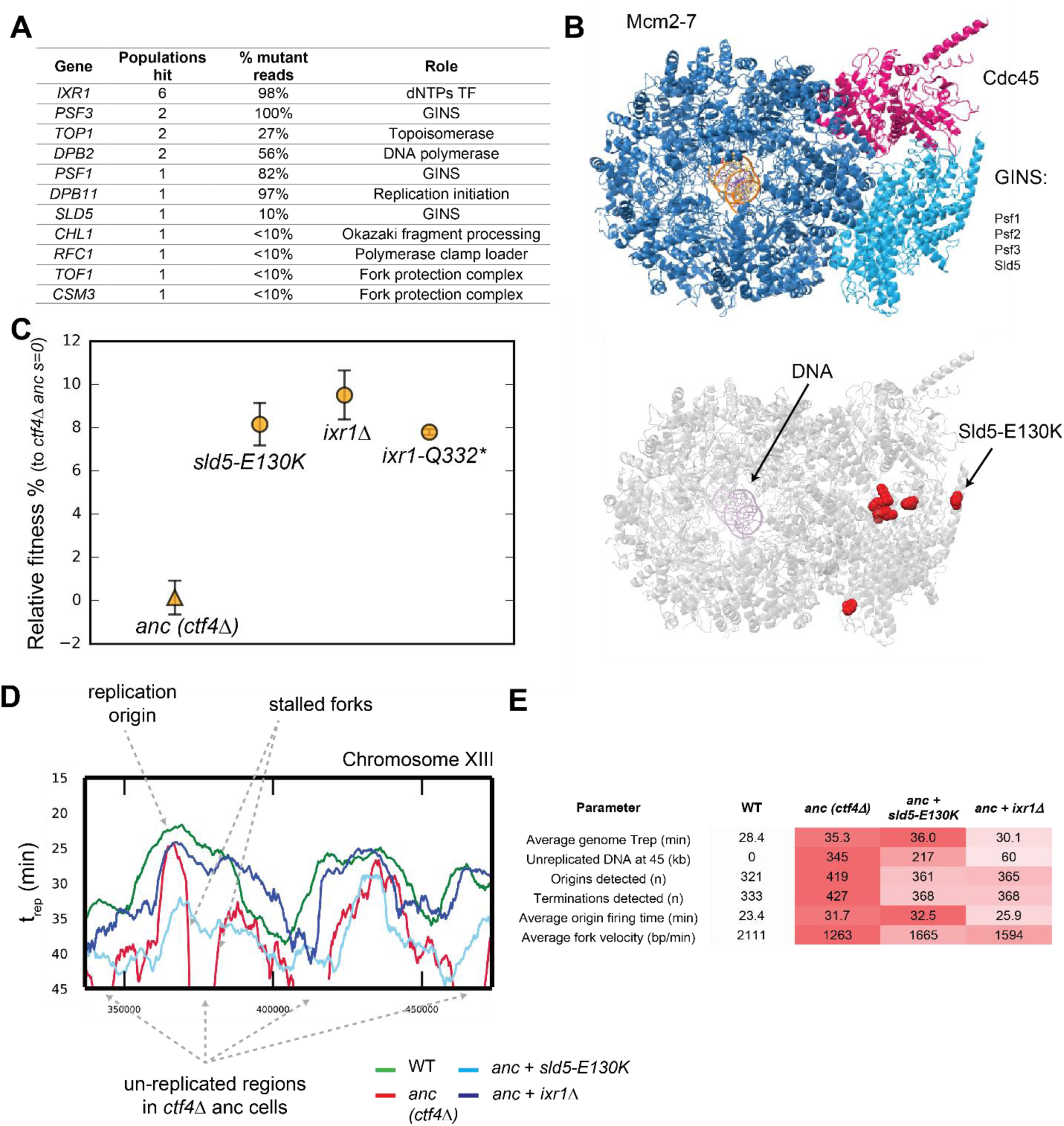
Adaptive mutations change DNA replication dynamics. **(A)** Genes involved in DNA replication that were mutated in evolved clones, and their role in replication. ‘populations hit’ is the number of populations where the gene was found mutated. ‘% Mutant reads’ was calculated as the average of the mutant read frequencies in the different populations where the mutation was detected. **(B)** Structure of the CMG helicase (PDB:5u8s, upper panel) highlighting the catalytic subunits (Mcm2-7) and the regulatory subunits (Cdc45 and GINS). Red spheres represent the residues affected by mutations found in evolved clones (lower panel). **(C)** The fitness of *ctf4*Δ strains carrying reconstructed mutations in the replicative helicase (*sld5-E129K*) and in *IXR1* (*ixr1*Δ and *ixr1-Q332*\*) relative to the *ctf4*Δ ancestor (*s*=0). Error bars represent standard deviations. **(D)** DNA replication profiles: cells were arrested in G1 and released into a synchronous S-phase, taking samples every 15 min for whole genome sequencing analysis. Change in DNA copy number over time were analyzed and used to calculate trep (time at which 50% of the cells in the population have replicated a given region (Figure S4C, see material and methods for details). A snapshot of chromosome XIII is shown as example, highlighting a DNA replication origin, the presence of stalled forks and unreplicated regions in *ctf4Δ* cells (which are absent in strains that also carry *sld5-E130K* or *ixr1*Δ mutations). **(E)** Quantitative analysis of DNA replication. Each parameter was derived from the genome-wide DNA replication profile of each sample (Fig. S4a, see material and methods for details). Heatmaps refer to the severity of the defect (white = wt, red = ctf4Δ ancestor).

*IXR1*, a gene indirectly linked to DNA replication, was mutated in several populations (Figure 4A). *IXR1* encodes for a transcription factor that indirectly and positively regulates the concentration of deoxyribonucleotide triphosphates (dNTPs, Tsaponina et al., 2011), the precursors for DNA synthesis. The occurrence of multiple nonsense mutations in this gene strongly suggested selection to inactivate Ixr1 (Figure S4D). Consistent with this prediction, we found that engineering either a nonsense mutation (*ixr1-Q332*\*) or a gene deletion conferred a selective advantage to *ctf4*Δ ancestor cells (Figure 4C).

We asked how mutations in the replicative helicase or inactivation of *IXR1* increased the fitness of *ctf4*Δ cells. One hypothesis is that the absence of Ctf4 reduces coordination of the activities required to replicate DNA and leads to the appearance of large regions of single stranded DNA, which in turn exposes the forks to the risk of nuclease cleavage or collapse. If this were true, slowing the replicative helicase or the synthesis of the leading strand would reduce the amount of single stranded DNA near the replication fork and improve the ability to complete DNA replication before cell division. To test this idea, we used whole genome sequencing at different points during a synchronous cell cycle to compare the dynamics of DNA replication in four strains: WT, the *ctf4Δ* ancestor, and *ctf4*Δ strains containing either the *sld5-E130K* or *ixr1Δ* mutations.

We found that cells lacking Ctf4 experience several defects compared to WT: on average, origins of replication fire later and DNA replication forks proceed more slowly across replicons, often showing fork stalling (Figure 4D, 4E, S4A). As a consequence of these two defects, cells still contain significant regions of unreplicated DNA late in S-phase (45 minutes, Figure 4D, 4E, S4a). Both *sld5-E130K* or *ixr1Δ* mutations significantly increase the average replication fork velocity primarily by avoiding stalls in DNA replication and thus leading to earlier replication of the regions that replicate late in the ancestral *ctf4*Δ cells (Figure 4D, 4E, S4A). Altogether, these results show that cells evolved modified DNA replication dynamics to compensate for defects induced by DNA replication stress.

### Epistatic interactions among adaptive mutations dictate evolutionary trajectories

Can we explain how the ancestral *ctf4*Δ strains recovered to within 10% of WT fitness in only 1000 generations? Although all the mutations that we engineered into *ctf4*Δ ancestor cells reduce the cost of DNA replication stress, none of them, individually, account for more than a third of the fitness increase observed over the course of the entire evolution experiment (Figure 1C). Sequencing individual evolved clones revealed the presence of mutations in at least two of the three modules whose effects we analyzed in isolation (Figure S5B, Table S1). We therefore asked if we could recapitulate the fitness of the evolved clones by adding adaptive mutations from multiple different modules to the *ctf4*Δ ancestor. We obtained all possible combinations of two, three, and four adaptive mutants, in the *ctf4*Δ ancestor, by sporulating a diploid strain that was heterozygous for all four classes of adaptive mutations: inactivation of the DNA damage checkpoint (*rad9Δ*), amplification of the cohesin loader (an extra copy of *SCC2*), alteration of the replicative helicase (*sld5-E130K*), and altered regulation of dNTP pools (*ixr1*Δ). We found that the two mutations that affected DNA replication were negatively epistatic (Figure 5A): in the presence of *ctf4*Δ, strains that contained both *sld5-E130K* and *ixr1Δ* were not significantly more fit than strains that contained only *ixr1Δ* and the quadruple mutant (*2X-SCC, rad9*Δ, *sld5-E130K, ixr1Δ*) was much less fit than the two triple mutants that contained only one of the two mutations that affected DNA replication (*2X-SCC, rad9Δ*, *sld5-E130K* and *2X-SCC, rad9Δ*, *ixr1Δ*). As a result, the two fittest strains carry only three mutations: in both cases, they affected the three modules we previously characterized: sister chromatid linkage and chromosome segregation *(2X-SCC2)*, the DNA damage checkpoint *(rad9Δ)* and DNA replication *(sld5-E130K* or *ixr1Δ)*. These two strains displayed a fitness comparable to the average of the evolved populations (Figure 1C), suggesting that we had recapitulated the major adaptive events in our engineered strains.

**Figure 5.**
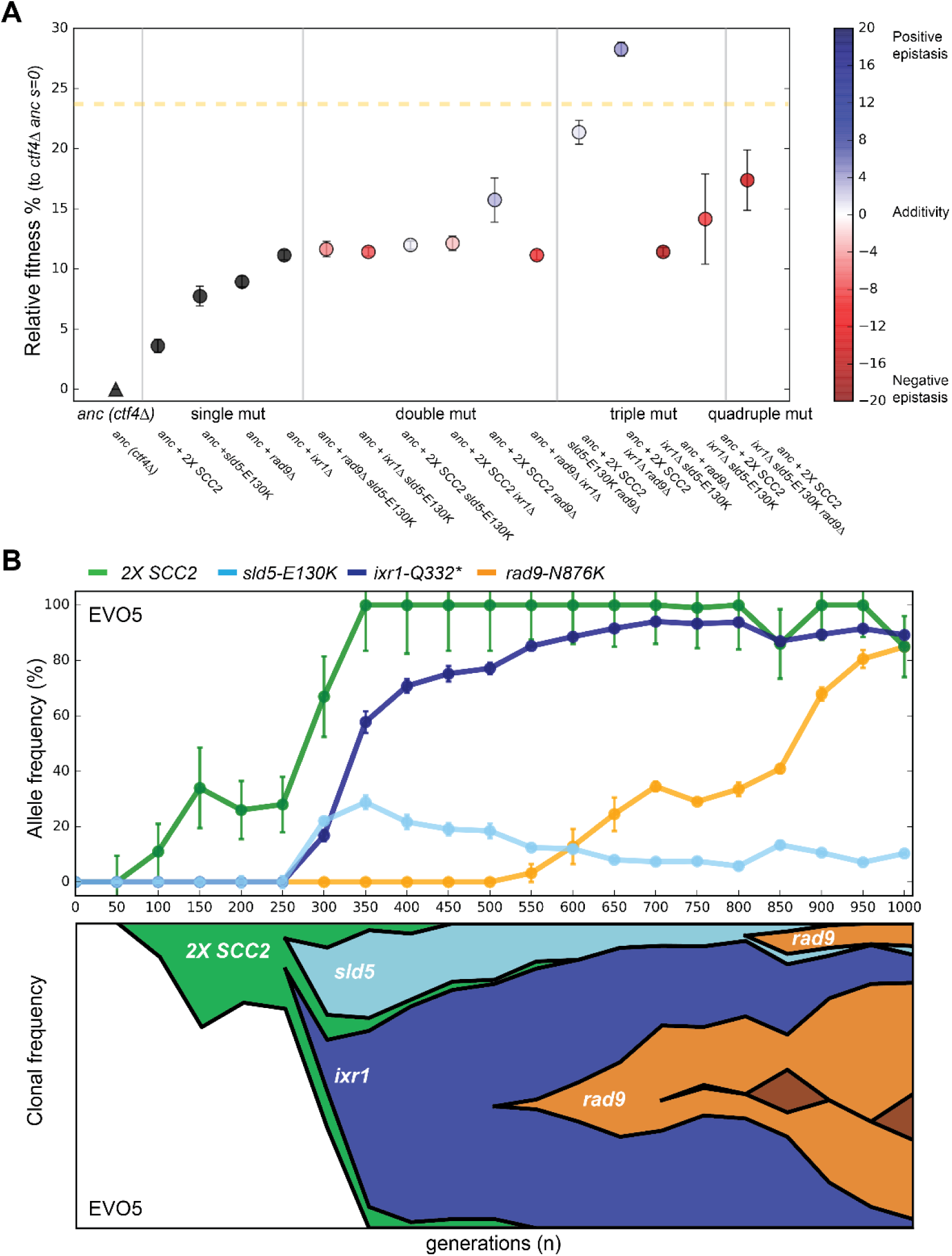
Epistatic interaction and evolutionary dynamics. **(A)** Fitness of all possible combinations of four adaptive mutations in the *ctf4Δ ;* ancestral background. The fitness measurements are relative to *ctf4Δ ;* ancestors (*s*=0). Dashed yellow line represents the average fitness of clones isolated from EVO5. Note that, differently from Figure 1C, fitness values are calculated relative to ancestors *ctf4Δ*, and not WT (hence the differences in absolute values, see material and methods). Error bars represent standard deviations. The fitnesses of individual strains are colored using the heatmap to the right of the figure, which represents epistasis: white = perfect additivity, red = negative epistasis (antagonism), blue = positive epistasis (synergy). Colors in heatmap represents the deviation in percentage between the observed fitness and the one calculated by adding the fitness effects of the individual mutations. **(B)** The temporal spread of mutant alleles during the experimental evolution of population EVO5 (upper panel). Error bars represent standard deviations. Genomic DNA was extracted from population samples, mutated loci were PCR amplified and Sanger sequencing was used to measure allele ratios (upper panel). A Muller diagram representing the lineages evolving in population EVO5 (lower panel). Data was obtained by combining alleles frequencies with their linkage as revealed by whole genome sequencing of clones isolated from EVO5 (Figure S5A and table S1).

We asked if the antagonistic interaction between *sld5-E130K* and *ixr1Δ* seen in our reconstructed strains had also occurred in our evolution experiment. We focused on an evolved population (EVO5) that carried all the mutations described above and analyzed the allele frequency in the intermediate samples collected across the evolution experiment. By following the frequency of alleles within the population and sequencing individual clones, we found that the mutations in the three modules happened in three consecutive selective waves: first, cells acquired an extra copy of the cohesin loader-encoding gene *SCC2*, second, *ixr1-Q332\** and *sld5-E130K* appeared, simultaneously, in two different lineages, and finally *rad9-N876K* appeared independently in the two lineages containing either *ixr1-Q332\** or *sld5-E130K* (Figure 5B). After their initial appearance, the two lineages containing *ixr1-Q332\** or *sld5-E130K* competed with each other for the remainder of the experiment. In this population, both of the final lineages accumulated mutations whose interaction was nearly additive or positively epistatic and avoided combinations that show strong negative epistasis (Figure 5A, S5A). Thus, although negative epistasis exists, selection finds trajectories that avoid it, as previously observed in a similar experiment perturbing cell polarity (Laan, Koschwanez, and Murray 2015).

## Discussion

Studying the molecular mechanism of evolutionary adaption helps to understand the balance between change and conservation during the evolution of biological functions. One approach is to compare processes in closely related organisms and use classical and molecular genetics to find the genetic variants responsible for inter-species differences. Another is to damage a process by applying a physiological stress that reduces the fitness of an organism and use experimental evolution to accumulate, identify and study the mutations that increase fitness and allow the organism to adapt to the stress.

We followed the latter approach and studied the evolutionary adaptation of cells experiencing constitutive DNA replication stress induced by the lack of a protein, Ctf4, that plays an important role in DNA replication. Over 1000 generations, populations increased from 75 to 90% of the fitness of their wild-type ancestors by sequentially accumulating mutations affecting three functions that contribute to chromosome metabolism: DNA replication, chromosome segregation and the cell-cycle checkpoint. We discuss the molecular mechanisms of adaptation, then consider how they interact to produce the final evolved phenotype, and close by commenting on the implications of our results for natural populations and cancer.

Cells lacking Ctf4 show an increased frequency of chromosome mis-segregation due to premature sister chromatid separation, but the mechanism underlying this defect is still unclear. Seven of our eight populations amplified *SCC2*, which encodes for one of the subunits of the cohesin loader complex. The simplest explanation for this result is that, the absence of Ctf4 restricts the productive loading of cohesin molecules that establish the linkage between sister chromatids. Amplifying the genes for the cohesin loader would increase its expression, increase the productive cohesin loading and improve the linkage between sister chromatids. Improving sister chromatid cohesion allows the evolved cells to segregate their chromosomes more accurately at mitosis, avoiding mitotic delays due to the spindle checkpoint, decreasing cell death and increasing fitness (Figure 6A).

**Figure 6.**
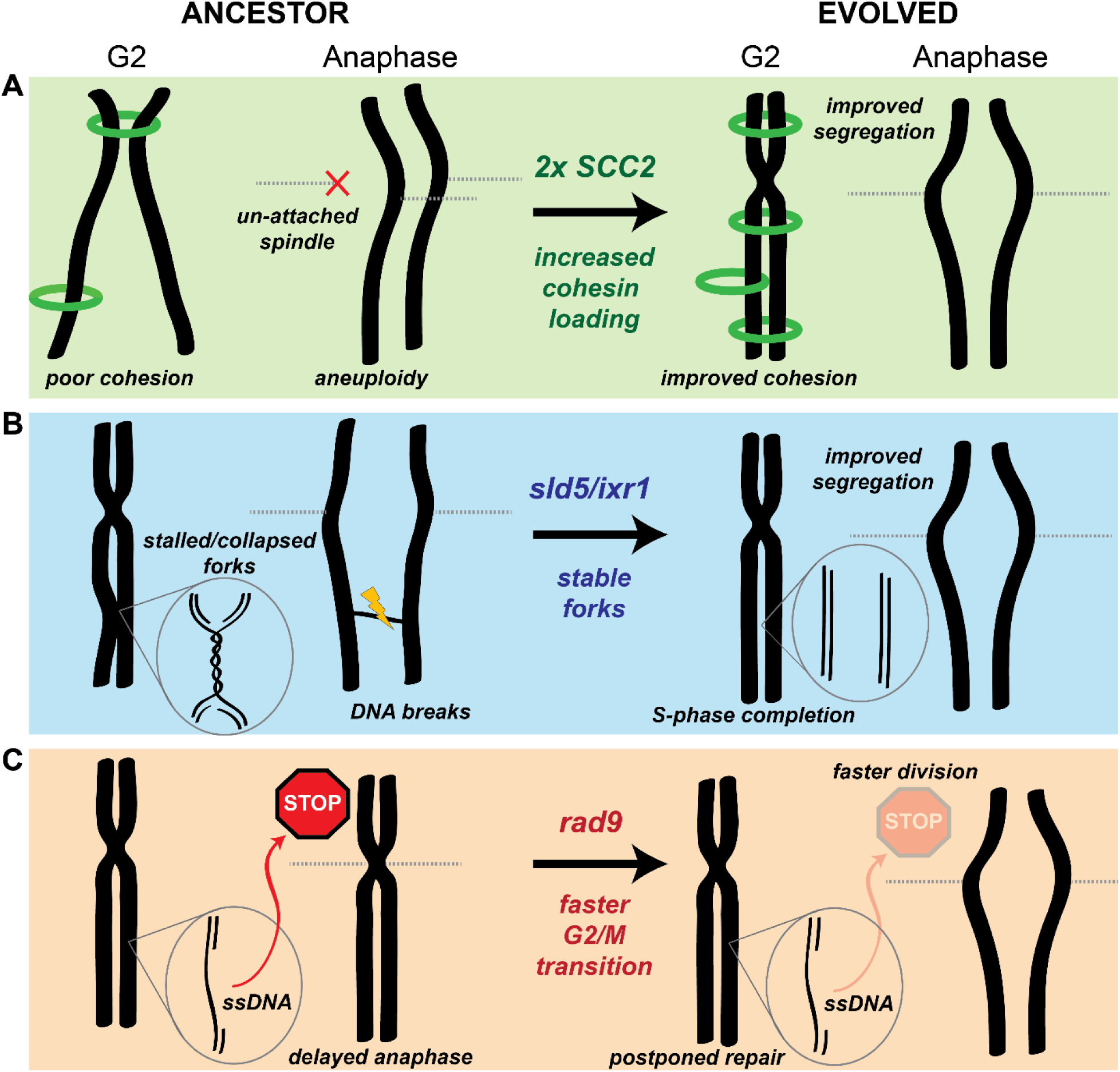
mechanistic models of adaptation. **(A)** Amplification of the cohesin loader subunit *SCC2* increases cohesin loading and sister chromatid cohesion leading to accurate chromosome segregation **(B)** Mutations of the replicative helicase (*sld5*) or in *ixr1* stabilize replication forks and ensure the completion of chromosome replication before anaphase. **(C)** mutations in *rad9* abolish the DNA damage checkpoint response triggered by stretches of single strand DNA (ssDNA) and allow faster cell division.

Persistent, cohesin-independent linkages between sister chromatids are an alternative source of segregation errors. These links include unreplicated regions of DNA or unresolved recombination structures (K.-L. Chan, North, and Hickson 2007; Ait Saada et al. 2017). If they persist after the removal of cohesin, they become lingering physical links (anaphase bridges) between sister chromatids that can lead to chromosome breakage or mis-segregation during anaphase (Gisselsson et al. 2000; K. L. Chan et al. 2009). Avoiding these problems requires that replication origins fire efficiently and replication forks move continuously. Our analysis of the dynamics of DNA replication argues that a combination of frequent fork stalling and slower origin firing causes under-replication of certain chromosomal regions in the ancestral *ctf4*Δ cells. We propose that these defects selected for mutations that have the apparently paradoxical effect of accelerating DNA replication by slowing down the replication forks: mutations like *sld5-E130K* and *ixr1*Δ make forks go slower and this reduced velocity stabilizes the forks, preventing frequent fork stalling or collapse and producing a higher overall fork velocity (Figure 6B). This hypothesis is consistent with two observations: first, although the *sld5* mutation is beneficial in ancestor cells, it decreases the fitness of WT cells (Figure S4E), a result we would expect from a slower replicative helicase. Second, reduced dNTPs concentrations reduce fork speed by slowing polymerase incorporation rates (Poli et al. 2012; Pai and Kearsey 2017; Koren, Soifer, and Barkai 2010) and inactivating Ixr1 reduces dNTP concentrations (Tsaponina et al. 2011). We tested this prediction by using an experimental system to manipulate dNTP concentrations: decreasing dNTP concentrations increased the fitness of *ctf4*Δ cells, while inducing higher dNTP production reduced fitness (Figure S4F).

Our evolved populations also accumulated mutations that inactivated the DNA damage checkpoint (Figure 2B-D). The benefit of these mutations arises from the loss of the DNA damage checkpoint‘s ability to delay the start of anaphase (Figure 2E, S2C). The absence of Ctf4 induces aberrant DNA structures and ssDNA that induce moderate activation of the checkpoint (Poli et al. 2012), which delays the start of anaphase, increasing doubling time and thus decreasing fitness (Figure 2D-E). Inactivating Rad9 eliminates the delay, shortening the time required for mitosis and increasing fitness (Figure 2E, S2C, 6C). This solution seems counter-intuitive, as the loss of a safeguard mechanism such as the DNA damage checkpoint should cause genetic instability in cells suffering from replication stress. The resolution of this paradox may lie in the overlapping action of the replication, DNA damage, and spindle checkpoints. We propose that the replication and the spindle checkpoints delay the cell cycle in response to defects that would kill the ancestral *ctf4*Δ cells, such as excessive replication fork collapses and pairs of sister chromatids attached to the same spindle pole, whereas the damage checkpoint responds to defects, like regions of single-stranded DNA, that can be repaired after cell division.

We asked how the mutants we identified and analyzed interacted with each other and whether they could explain the fitness of our evolved populations. Measuring allele frequencies over time and engineering all possible combinations of adaptive mutations allowed us to propose a detailed model for the evolutionary trajectories of our population 5 (EVO5). Segmental amplifications form at a higher frequency than other types of mutation (Lynch et al. 2008; Sharp et al. 2018; Yona, Frumkin, and Pilpel 2015); although most are detrimental, the amplification of specific genes can be advantageous and cause rapid adaptation (Gresham et al. 2008; Adamo et al. 2012; Hughes et al. 2000; Payen et al. 2014). Thus, the first event in EVO5 is the spread of a segmental amplification of chromosome IV containing *SCC2*, which improves fitness by reducing cohesion defects. In this lineage, mutations in the replicative helicase, *sld5-E130K*, and *ixr1-Q332\** were then detected almost simultaneously but in different clones. Above, we argue that both mutations slow replication forks. If there is an optimal fork speed in ctf4Δ cells, the presence of a second mutation of this class might be ineffective or even detrimental if the forks move too slowly, explaining the negative epistasis we observed. Because the *ixr1* and *sld5* mutations improve DNA replication to a similar extent, the two lineages have comparable fitness, explaining the clonal interference that persists for the rest of the experiment. The last mutation in EVO5 is an identical frameshift mutation in the two lineages that inactivates Rad9. Interestingly, loss of function mutations in *RAD9*, despite the large target size of this gene, only appear relatively late during the experiment (Figure 5A and S5B). Furthermore, they happen after other mutations have reduced some of the problems imposed by replication stress. This order suggests that a sustainable fitness advantage of mutations of the DNA damage checkpoint may depend on previous changes in the replication forks stability.

Overall, this study reveals the short-term evolutionary plasticity of chromosome metabolism. A single genetic perturbation that induces DNA replication stress and a thousand generations are enough to select for significant changes in modules affecting chromosome metabolism. By the end of the experiment, evolved lineages have sequentially modified chromosome cohesion, changed the speed of replication forks, and lost an important cell-cycle response to damage. Changes in these conserved modules collectively contribute to the evolved phenotype and allow cells to achieve high fitness despite the presence of constitutive DNA replication stress. This result suggests that despite their conservation, these modules are evolutionarily plastic and can accommodate short-term responses to strong perturbations, helping to explain differences that have accumulated over hundreds to billions of years of evolution.

### Implications for species evolution in the wild

Despite being conserved across much of evolution, some of the modules that collectively perform chromosome metabolism and maintain genomes show major important differences between clades, even within the eukaryotic kingdom (Gourguechon, Holt, and Cande 2013; Akiyoshi and Gull 2014; Y. Liu, Richards, and Aves 2009). For instance, a recent study found species in the yeast genus *Hanseniaspora* that lack several important genes implicated in cell cycle progress and DNA repair, including checkpoint factors such as *RAD9* and *MAD2* (Steenwyk et al. 2019). Trying to explain these differences is puzzling, especially if *ad-hoc* selectionist hypotheses are invoked for each different feature. For instance, what could select for a lack of an important safeguard such as the DNA damage checkpoint? The evolutionary plasticity of chromosome metabolism that we reveal in this work may help to explain differences like these: mutations in ancestor cells could initiate an evolutionary trajectory that progressively modifies modules that are functionally linked and ultimately leads to increased fitness. But what are the initial perturbations that trigger such changes in fundamental aspects of cell biology? The *ctf4*Δ cells that we evolved have a 25% fitness difference relative to their wild type ancestors, meaning that they would rapidly be eliminated from any population of reasonable size. Given the evolutionary rarity of major rearrangements in cell biology we can invoke events that are improbable including passing through very small populations bottlenecks or being attacked by selfish genetic elements whose molecular biology targets an important protein in an essential process. If the processes that were damaged during these events, were part of chromosome metabolism, the consequent evolutionary adaptation could lead to changes in the rates at which the structures of genomes evolve. An increase in these rates, in turn, could potentially accelerating speciation by making it easier for populations to acquire meiotically incompatible chromosome configurations.

### Implications for cancer evolution

Remarkably, our experiment recapitulates several phenomena observed during cancer development. Replication stress is thought to be a ubiquitous feature of cancer cells (Macheret and Halazonetis 2015) with oncogene activation leading to replication stress and genetic instability (Di Micco et al. 2006; Bartkova et al. 2006; Neelsen et al. 2013). The absence of Ctf4 in our ancestor cells causes several phenotypes observed in oncogene-induced DNA replication stress including late-replicating regions, elevated mutation rates, and chromosome instability (Muñoz and Méndez 2016; Macheret and Halazonetis 2015; Fumasoni et al. 2015). Furthermore, simply by propagating cells, we generated evolved lines that mimic many features seen in tumors: a) individual final populations contain genetically heterogeneous clones, often with different karyotypes characterized by aneuploidies and chromosomal rearrangements (Lengauer, Kinzler, and Vogelstein 1998; Laughney et al. 2015; Davoli et al. 2013), b) evolved lineages display altered DNA replication profiles compared both to WT cells and their mutant ancestors (Donley and Thayer 2013; Amiel et al. 1999), c) several lines have inactivated the DNA damage checkpoint (Schultz et al. 2000; Hollstein et al. 1991), and d) improved sister chromatid cohesion (Rhodes, McEwan, and Horsfield 2011; Sarogni et al. 2019; Xu et al. 2011). All these features are adaptive in our populations, suggesting that similar changes in cancer cells may be the result of selection and contribute to the accumulation of other cancer hallmarks during cancer evolution. The similarities between tumorigenesis and our experiment lead us to speculate that a major selective force in the early stages of tumor evolution is the need to counteract the fitness costs of replication stress. Understanding the evolutionary mechanisms and dynamics of the adaptation to replication stress could therefore shed light on the early stage of tumor development.

### Perspective

In this work, we identified the main adaptive strategies that cells use to adapt to DNA replication stress induced by the absence of Ctf4. Our results reveal that defects in one function can be compensated for by two types of mutations: those in the original function and those in functions that are biologically coupled to it. Focusing on less common adaptive strategies, apparently unlinked to chromosome metabolism, could therefore potentially identify novel players that affect genome stability. It would also be interesting to induce DNA replication stress by other means, such as de-regulating replication initiation or by inducing re-replication. Analyzing the response to these challenges will reveal whether the DNA replication module has a common or diverse set of evolutionary strategies to different perturbations. Finally, this approach could be extended to many other types of cellular stress, potentially revealing other molecular adaption aspects that could collectively help understanding cellular evolution.

## Material and methods

### Strains

All strains were derivatives of a modified version (Rad5^+^) of *S. cerevisiae* strain W303 (*leu2-3,112 trp1-1 can1-100 ura3-1 ade2-1 his3-11,15, RAD5*+). TableS4 lists each strain‘s genotype. The ancestors of WT and *ctf4Δ* strains were obtained by sporulating a *CTF4/ctf4Δ* heterozygous diploid. This was done to minimize the selection acting on the ancestor strains before the beginning of the experiment. Diploid stains were grown on YPD, transferred to sporulation plates (sodium acetate 0.82%, potassium chloride 0.19%, sodium chloride 0.12%, magnesium sulfate 0.035%) and incubated for four days at 25°C. Tetrads were re-suspended in water containing zymolyase (Zymo research, 0.025 u/μl), incubated at 37°C for 45 sec, and dissected on a YPD plate using a Nikon eclipse E400 microscope equipped with a Schuett-Biotec TDM micro-manipulator. Spores were allowed to grow into visible colonies and genotyped by presence of genetic markers and PCR.

### Media and growth conditions

Standard rich media, YPD (1% Yeast-Extract, 2% Peptone, 2% D-Glucose) was used for all experiments except in the experiment in Figure S4F where YP + 2% raffinose and YP + 2% raffinose + 2% galactose were also used. Cells were synchronized either in metaphase, for 3hrs in YPD containing nocodazole (8 μg/ml, in 1% DMSO) or in G1, for 2hrs in YPD, pH 3.5 containing α-factor (3 μg/ml). Synchronization was verified by looking at cell morphology. In the experiment in Figure 2E, cells were then washed twice in YPD containing 50 μg/ml pronase and released in S-phase at 30°C in YPD. α-factor (3 μg/ml) was added again at 30 min to prevent a second cell cycle from occurring.

### Experimental evolution

The 16 populations used for the evolution experiment were inoculated in glass tubes containing 10ml of YPD from 8 *ctf4Δ* colonies (EVO1-8) and 8 WT colonies (EVO9-16). All the colonies were derived by streaking out *MAT****a*** (EVO1-4 and EVO9-12) or *MATα* (EVO5-9 and EVO13-16) ancestors. Glass tubes were placed in roller drums at 30°C and grown for 24hrs. Daily passages were done by diluting 10 μl of the previous culture into 10 ml of fresh YPD (1:1000 dilution, allowing for approximately 10 generations/cycle). All populations were passaged for a total of 100 cycles (≈1000 generations). Every 5 cycles (≈50 generations) 800 μl of each evolving population was mixed with 800 μl of 30% v/v glycerol and stored at -80°C for future analysis (Figure 1B). After 1000 generations four evolved clones were isolated from the each of the eight *ctf4*Δ evolved populations (a total of 32 clones) by streaking cells on a YPD plate. Single colonies were then grown in YPD media and saved in glycerol at -80°C as for the rest of the samples.

### Whole genome sequencing

Genomic DNA library preparation was performed as in (Koschwanez, Foster, and Murray 2013) with an Illumina Truseq DNA kit. Libraries were then pooled and sequenced either with an Illumina HiSeq 2500 (125bp paired end reads) or an Illumina NovaSeq (150bp paired end reads). The samtools software package (samtools.sourceforge.net) was then used to sort and index the mapped reads into a BAM file. GATK (www.broadinstitute.org/gatk, McKenna et al., 2010) was used to realign local indels, and Varscan (varscan.sourceforge.net) was used to call variants. Mutations were found using a custom pipeline written in Python (www.python.org). The pipeline (github.com/koschwanez/mutantanalysis) compares variants between the reference strain, the ancestor strain, and the evolved strains. A variant that occurs between the ancestor and an evolved strain is labeled as a mutation if it either (1) causes a non-synonymous substitution in a coding sequence or (2) occurs in a regulatory region, defined as the 500 bp upstream and downstream of the coding sequence (Table S1).

### Identification of putative adaptive mutations

Three complementary approaches were combined to identify the putative modules and genes targeted by selection.

#### Convergent evolution on genes

This method relies on the assumption that those genes that have been mutated significantly more than expected by chance alone, represent cases of convergent evolution among independent lines. The mutations affecting those genes are therefore considered putatively adaptive. The same procedure was used independently on the mutations found in WT and *ctf4Δ* evolved lines:

We first calculated per-base mutation rates as the total number of mutations in coding regions occurring in a given background (*ctf4*Δ evolved or WT evolved), divided by the size of the coding yeast genome in bp (including 1000bp per ORF to account for regulatory regions)

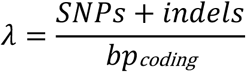

If the mutations were distributed randomly in the genome at a rate λ, the probability of finding n mutations in a given gene of length *N* is given by the Poisson distribution:

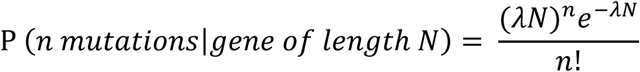

For each gene of length *N*, we then calculated the probability of finding ≥ n mutations if these were occurring randomly.

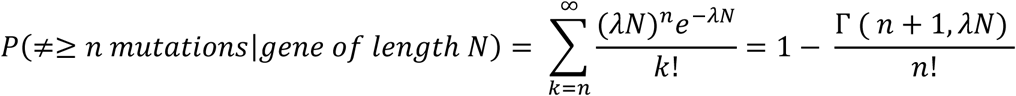

(Where Γ is the upper incomplete gamma function) which gives us the p-value for the comparison of the observed mutations with the null, Poisson model. In order to decrease the number of false positives, we then performed multiple-comparison corrections. The more stringent Bonferroni correction (α=0.05) was applied on the WT evolved mutations dataset, while Benjamini-Hochberg correction (α=0.05) was used for the *ctf4Δ* mutation dataset. Genes that were found significantly selected in the evolved WT clones (after Bonferroni correction) were removed from the list of evolved *ctf4Δ* strains. This is because, since they were target of selection even in WT cells, they are likely involved in processes that are un-related to DNA replication and are instead associated with adaptation to sustained growth by serial dilutions. TableS2 lists the mutations detected in evolved *ctf4Δ* clones, after filtering out those that occurred in genes that were significantly mutated in the WT populations. Genes significantly selected in these clones are shown in dark grey (after Benjamini-Hochberg correction with α=0.05).

#### Bulk segregant analysis

Bulk segregants analysis experimentally identifies putative adaptive mutations present in a given evolved clone. Briefly, a clone is selected from the population and then backcrossed to a derivative of the WT ancestor. The resulting diploid is sporulated, allowing the mutant alleles accumulated during 1000 generations to randomly segregate among the haploid progeny. The haploid progeny is then selected for growth (and for *ctf4Δ*) for 50-80 generations in rich media. This regime, as in the experimental evolution, selects for cells with higher fitness. The cells with causal alleles therefore quickly increase their frequency within the selected population. Non-causal alleles segregate randomly and, since they don‘t contribute to fitness, they are expected to be present in half of the cells at the end of the progeny selection. Deep sequencing of the genomic DNA extracted from the selected progeny population reveal the alleles frequencies and allows the identification of the ones that segregate with the evolved phenotype (frequency >70% in our case). Bulk segregant analysis was adapted from (Koschwanez, Foster, and Murray 2013). One clone per population was selected for further analysis (Figure S1B). In these clones, the original *ctf4Δ* genetic marker *ble* was substituted with a *KanMX6* cassette by homologous recombination, to allow for a more efficient selection. *ura3-1* evolved clones were mated with either a *MAT****a*** or *MAT*α, *ura3∷NatMX4-pSTE2-URA3* derivative of the WT ancestor. In this strain, the endogenous *URA3* promoter is replaced with the *STE2* promoter, which is only induced in *MAT****a*** cells, making it possible to select for *MAT****a*** spores after meiosis. Mating was performed by mixing cells from the two strains together on a YPD plate with a toothpick and growing overnight at 30°C. The mating mixtures were then plated on double selective media, and a diploid strain from each cross was selected from a colony on the plate. To sporulate the diploid strains, cultures were grown to saturation in YPD, and then diluted 1:100 into YP 2% acetate. The cells were grown in acetate for 12 hrs, pelleted and resuspended in 2% acetate. After 5 days of incubation on a roller drum at 25°C, sporulation was verified by observing the formation of tetrads under the microscope. To digest ascii, 10 ml of the sporulated culture was pelleted and resuspended in 500 μl with 250 units of Zymolyase for 1 hr at 30°C. 4000 μl of water and 500 μl of 10% Triton X-100 were added, and the digested spores were then sonicated for 1 min to separate the tetrads. The spores were spun down slowly (6000 rpm) and resuspended in 50 ml of -URA +G418 media. This media selects for *ctf4Δ* haploid *MAT****a*** cells: neither haploid *MAT*α nor diploid *MAT****a***/*MAT*α cells can express *URA3* from the *STE2* promoter. Each culture was then diluted 1:100 in fresh -URA + G418 media for 10 consecutive passages, allowing for ≈66 generations to occur. Genomic DNA was extracted from the final saturated culture and used for library preparation and whole genome sequencing as described.

#### Convergent evolution on modules

Statistical methods to find frequently mutated genes are focused on individual genes that contribute to an evolved trait. Functions that can be modified by affecting several genes would be therefore under-represented in the previous analysis. To account for this, we looked for gene ontology (GO) terms enriched among the mutations found to be positively selected in *ctf4Δ* evolved clones (tableS1, dark rows), or found segregating with the evolved phenotype by bulk segregant analysis (Figure S1B). The combined list of mutations was input as ‘multiple proteins’ in the STRING database, which reports on the network of interactions between the input genes (https://string-db.org). Several GO terms describing pathways involved in the DNA and chromosome metabolism were found enriched among the putative adaptive mutations provided (Figure S1C and Table S3). Since GO terms are often loosely defined and partially overlapping, we manually identified, based on literature search, four modules as putative targets of selection: DNA replication, chromosome segregation, cell cycle checkpoints, and chromatin modifiers. The full list of mutated genes observed in the evolved *ctf4Δ* clones was then used as input in the STRING database. This was done to account for genes, that despite not being identified as containing adaptive mutations by the previous techniques, are part of modules under selection: mutations in these genes could have contributed to the final phenotype. The interaction network between mutated genes was downloaded and curated in Cytoscape. For clarity of representation, only those nodes strongly connected to the previously identified modules are shown in Figure 1D.

### Fitness assays

To measure relative fitness, we competed the ancestors and evolved strains against reference strains. Both WT (Figure 1C, S1A, S4E, S5A) and *ctf4Δ* (Figure 2D, S2C, 3C, 4C, S4E-F, 5A) reference strains were used. A *pFA6a-prACT1-yCerulean-HphMX4* plasmid was digested with *Age*I and integrated at one of the *ACT1* loci of the original heterozygous diploid (*CTF4/ctf4Δ*) strain. This allow for the expression of fluorescent protein yCerulean under the strong actin promoter. The heterozygous diploid was then sporulated and dissected to obtain fluorescent WT or *ctf4Δ* reference haploid strains. For measuring the relative fitness, 10 ml of YPD were inoculated in individual glass tubes with either the frozen reference or test strains. After 24 hrs the strains were mixed in fresh 10 ml YPD tubes at a ratio dependent on the expected fitness of the test strain compared to the reference (i.e 1:1 if believed to be nearly equally fit) and allowed to proliferate at 30°C for 24 hrs. 10 μl of samples were taken from this mixed culture (day 0) and the ratio of the two starting strains was immediately measured. Tubes were then cultured following in the same conditions as the evolution experiment by diluting them 1:1000 into fresh media every 24hrs for 4 days, monitoring the strain ratio at every passage. Strain ratios and number of generations occurred between samples were measured by flow cytometer (Fortessa, BD Bioscience). Ratios *r* were calculated based on the number of fluorescent and non-fluorescent events detected by the flow cytometer:

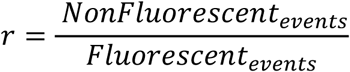

Generations between time points *g* were calculated based on total events measured at time 0 hr and time 24 hrs:

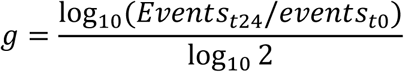

Linear regression was performed between the (g, log_*e*_ *r*) points relative to every sample. Relative fitness *s* was calculated as the slope of the resulting line. Note that the absolute values of relative fitness change depending on the reference strain used: a strain that shows 27% increased fitness when measured against *ctf4Δ* (that is 27% less fit then WT), does not equate the WT fitness. This is because a 27% increase of 0.73 (*ctf4Δ* fitness compared to WT) gives 0.93, hence a 7% fitness defect compared to WT.

### Cell cycle profiles

Cell cycle analysis was conducted as previously described (Fumasoni et al. 2015). In brief, 1×10^7^ cells were collected from cultures by centrifugation, and resuspended in 70% ethanol for 1 hr. Cells were then washed in 50 mM Tris-HCl (pH 7.5), resuspended in the same buffer containing 0.4 μg/ml of RNaseA and incubated at 37°C for at least 2 hrs. Cells were collected and further treated overnight at 37°C in 50 mM Tris-HCl (pH 7.5) containing proteinase K (0.4 μg/ml). Cells were then centrifuged and washed in 50 mM Tris-HCl (pH 7.5). Samples were then diluted 10-20-fold in 50 mM Tris-HCl (pH 7.8) containing 1 mM Sytox green, and analyzed by flow cytometer (Fortessa, BD Bioscience). The FITC channel was used to quantify the amounts of stained-DNA per cell. Cell cycle profiles were analyzed and visualized in Flowjo (BD). The percentage of genome replicated at 30 min was calculated based on the cell cycle profile as follow *G*_*rep*_ = *DNA content mode*/(2*C* – 1*C*) * 100. The height of the 1C and 2C peaks was obtained as the max cells count reached by the respective peak.

### Copy number variations (CNVs) detection by sequencing

Whole genome sequencing and read mapping was done as previously described. The read-depths for every unique 100 bp region in the genome were then obtained by using the VarScan copynumber tool. A custom pipeline written in python was used to visualize the genome-wide CNVs. First, the read-depths of individual 100 bp windows were normalized to the genome-wide median read-depth to control for differences in sequencing depths between samples. The coverage of the ancestor strains was then subtracted from the one of the evolved lines to reduce the noise in read depth visualization due to the repeated sequences across the genome. The resulting CNVs were smoothed across five 100 bp windows for a simpler visualization. Final CNVs were then plotted relative to their genomic coordinate at the center of the smoothed window. Since the WT CNVs were subtracted from the evolved CNVs, the y axis refers to the copy number change occurred during evolution (i.e. +1 means that one an extra copy of a chromosome fragment has been gained).

### Premature sister chromatid separation assay

Logarithmically growing cells were arrested in metaphase as previously described. Samples were then collected and fixed in 4% formaldehyde for 5 min at room temperature. Cells were washed In SK buffer (1M sorbitol, 0.05 M K2PO4) and sonicated for 8 seconds prior to microscope analysis. Images were acquired with a Nikon eclipse Ti spinning-disk confocal microscope using a 100X oil immersion lens. Fluorescence was visualized with a conventional FITC excitation filter and a long pass emission filter. Images were analyzed using ImageJ.

### DNA replication profiles

DNA replication profiling was adapted from Müller et al. 2014; Saayman, Ramos-Pérez, and Brown 2018; Bar-Ziv, Voichek, and Barkai 2016. Genomic DNA, library preparation and CNVs detection were performed independently on all the collected samples as previously described. A custom python script was used to analyze the CNVs from multiple time points from the same strain to produce DNA replication profiles. Read-depths of individual 100 bp windows were normalized to the genome-wide median read-depth to control for differences in sequencing depths between consecutive samples. To allow for intra-strain comparison, coverage was then scaled according to the sample DNA content measured as the median of the cell-cycle profile obtained by flow cytometry. The resulting coverage was then averaged across multiple 100 bp windows and a polynomial data smoothing filter (Savitsky-Golay) was applied to the individual coverage profiles to filter out noise. Replication timing t_rep_ is defined as the time at which 50% of the cells in the population replicated a given region of the genome (Figure S4C), which is equivalent to an overall relative coverage of 1.5x, since 1x corresponds to an unreplicated region and 2x to a fully replicated one. The replication timing t_rep_ was calculated by linearly interpolating the two time points with coverage lower and higher than 1.5x and using such interpolation to compute the time corresponding to 1.5x coverage. Final t_rep_ were then plotted relative to their window genomic coordinates. Unreplicated regions at 45 min were calculated as the sum of all regions with t_rep_>45min. To find DNA replication origins, the t_rep_ profiles along the genome were filtered using a Fourier low-pass filter to remove local minima and then used to find local peaks. Only origins giving rise to long replicons were used to measure fork velocity. Fork velocity was calculated by dividing the distance between the origin and the closest termination site by the time required to replicate the region. Duplicate replication profiles were obtained from two independent experiments for each strain. Reproducibility was confirmed with qualitatively and quantitatively comparable results across duplicates. The reliability of the pipeline was assessed by qualitatively and quantitatively comparing our WT results with previously reported measurements (Raghuraman et al. 2001; Müller et al. 2014).

### Analysis of allele frequency by sanger sequencing

Allele frequencies within populations were estimated as in (Koschwanez, Foster, and Murray 2013). In brief, chromatograms obtained by sanger sequencing were used to estimate the fraction of mutant alleles in a population at different time points during the evolution. The fraction of mutant alleles in the population was assumed to be the height of the mutant allele peak divided by the height of the mutant allele peak plus the ancestor allele peak. The values from two independent sanger sequencing reactions, obtained by primers lying upstream and downstream the mutations, were averaged to obtain the final ratios. Values below the approximate background level were assumed to be zero, and values above 95% were assumed to be 100%.

### Segmental amplification detection by digital PCR

Droplet digital PCR was used to detect the amplifications of the fragment containing *SCC2* at different time points during evolution. Genomic DNA was prepared and diluted accordingly. Bio-Rad ddPCR supermix for probes (no dUTP) was used to prepare probes specific to *SCC2* and the centromere of chromosome IV. A Bio-Rad QX200 Droplet Generator was used to generate droplets containing genomic DNA and probes. The droplet PCR was performed in a Bio-Rad thermocycler and analyzed with a Bio-Rad QX200 Droplet Reader. SCC2/Centromere ratios were then used to quantify SCC2 copy numbers. To estimate the percent of cells carrying the SCC2 amplification within a population we assumed that the allele spreading in the population was a duplication of SCC2 (as indicated by the EVO5 copy number analysis). Values above 95% were assumed to be 100%.

## Supporting information

Mutations list

Selected genes

Enriched GO terms

## Acknowledgments

We thank Stephen Elledge and Philip Zegerman for sharing yeast strains; Andrea Giometto, Mayra Garcia and John Koschwanez for assistance in data analysis; Stephen Bell, Michael Desai, Michael Laub, Bodo Stern, Sriram Srikant, Thomas LaBar and Yi Chen for critical reading of the manuscript; Claire Hartman and Zachary Niziolek from the Harvard Bauer Core Facility for technical assistance. Yoav Voichek and Felix Jonas for advice on DNA replication profiling; We thank the members of the Murray and Nelson labs for helpful discussions. This work was supported by NIH grant RO1-GM43987 and by the NSF-Simons Center for Mathematical and Statistical Analysis of Biology at Harvard (#1764269) to AWM. MF gratefully acknowledges fellowship support from the Human Frontiers Science Program (LT000786/2016-L), EMBO (ALTF 485-2015) and AIRC (iCARE 17957).

## Author contributions

MF designed and performed the research, analyzed and interpreted the data, and wrote the paper. AWM designed the research, interpreted the data, and wrote the paper.

## References

Abe, Takuya, Ryotaro Kawasumi, Michele Giannattasio, Sabrina Dusi, Yui Yoshimoto, Keiji Miyata, Koyuki Umemura, Kouji Hirota, and Dana Branzei. 2018. “AND-1 Fork Protection Function Prevents Fork Resection and Is Essential for Proliferation.” Nature Communications 9 (1): 3091. https://doi.org/10.1038/s41467-018-05586-7.

Adamo, G. M., M. Lotti, M. J. Tamas, and S. Brocca. 2012. “Amplification of the CUP1 Gene Is Associated with Evolution of Copper Tolerance in Saccharomyces Cerevisiae.” Microbiology 158 (Pt_9): 2325–35. https://doi.org/10.1099/mic.0.058024-0.

Ait Saada, Anissia, Ana Teixeira-Silva, Ismail Iraqui, Audrey Costes, Julien Hardy, Giulia Paoletti, Karine Fréon, and Sarah A.E. Lambert. 2017. “Unprotected Replication Forks Are Converted into Mitotic Sister Chromatid Bridges.” Molecular Cell 66 (3): 398-410.e4. https://doi.org/10.1016/j.molcel.2017.04.002.

Akiyoshi, Bungo, and Keith Gull. 2014. “Discovery of Unconventional Kinetochores in Kinetoplastids.” Cell 156 (6): 1247–58. https://doi.org/10.1016/J.CELL.2014.01.049.

Amiel, A., I. Kirgner, E. Gaber, Y. Manor, M. Fejgin, and M. Lishner. 1999. “Replication Pattern in Cancer: Asynchronous Replication in Multiple Myeloma and in Monoclonal Gammopathy.” Cancer Genetics and Cytogenetics 108 (1): 32–37. https://doi.org/10.1016/S0165-4608(98)00107-1.

Aves, Stephen J, Yuan Liu, and Thomas A Richards. 2012. “Evolutionary Diversification of Eukaryotic DNA Replication Machinery.” Sub-Cellular Biochemistry 62 (January): 19–35. https://doi.org/10.1007/978-94-007-4572-8_2.

Bar-Ziv, Raz, Yoav Voichek, and Naama Barkai. 2016. “Chromatin Dynamics during DNA Replication.” Genome Research 26 (9): 1245–56. https://doi.org/10.1101/gr.201244.115.

Barrick, Jeffrey E., and Richard E. Lenski. 2013. “Genome Dynamics during Experimental Evolution.” Nature Reviews Genetics 14 (12): 827–39. https://doi.org/10.1038/nrg3564.

Bartkova, Jirina, Nousin Rezaei, Michalis Liontos, Panagiotis Karakaidos, Dimitris Kletsas, Natalia Issaeva, Leandros-Vassilios F. Vassiliou, et al. 2006. “Oncogene-Induced Senescence Is Part of the Tumorigenesis Barrier Imposed by DNA Damage Checkpoints.” Nature 444 (7119): 633–37. https://doi.org/10.1038/nature05268.

Bell, Stephen P., and Karim Labib. 2016. “Chromosome Duplication in Saccharomyces Cerevisiae.” Genetics 203 (3).

Branzei, Dana, and Marco Foiani. 2010. “Maintaining Genome Stability at the Replication Fork.” Nature Reviews. Molecular Cell Biology 11 (MARCh): 208–19. https://doi.org/10.1038/nrm2852.

Burhans, William C, and Martin Weinberger. 2007. “DNA Replication Stress, Genome Instability and Aging.” Nucleic Acids Research 35 (22): 7545–56. https://doi.org/10.1093/nar/gkm1059.

Buskirk, Sean W, Ryan Emily Peace, and Gregory I Lang. 2017. “Hitchhiking and Epistasis Give Rise to Cohort Dynamics in Adapting Populations.” Proceedings of the National Academy of Sciences of the United States of America 114 (31): 8330–35. https://doi.org/10.1073/pnas.1702314114.

Chan, Kok-Lung, Phillip S North, and Ian D Hickson. 2007. “BLM Is Required for Faithful Chromosome Segregation and Its Localization Defines a Class of Ultrafine Anaphase Bridges.” The EMBO Journal 26 (14): 3397–3409. https://doi.org/10.1038/sj.emboj.7601777.

Chan, Kok Lung, Timea Palmai-Pallag, Songmin Ying, and Ian D. Hickson. 2009. “Replication Stress Induces Sister-Chromatid Bridging at Fragile Site Loci in Mitosis.” Nature Cell Biology 11 (6): 753–60. https://doi.org/10.1038/ncb1882.

Ciosk, Rafal, Masaki Shirayama, Anna Shevchenko, Tomoyuki Tanaka, Attila Toth, Andrej Shevchenko, and Kim Nasmyth. 2000. “Cohesin’s Binding to Chromosomes Depends on a Separate Complex Consisting of Scc2 and Scc4 Proteins.” Molecular Cell 5 (2): 243–54. https://doi.org/10.1016/S1097-2765(00)80420-7.

Cross, Frederick R, Nicolas E Buchler, and Jan M Skotheim. 2011. “Evolution of Networks and Sequences in Eukaryotic Cell Cycle Control.” Philosophical Transactions of the Royal Society of London. Series B, Biological Sciences 366 (1584): 3532–44. https://doi.org/10.1098/rstb.2011.0078.

Davoli, Teresa, Andrew Wei Xu, Kristen E. Mengwasser, Laura M. Sack, John C. Yoon, Peter J. Park, and Stephen J. Elledge. 2013. “Cumulative Haploinsufficiency and Triplosensitivity Drive Aneuploidy Patterns and Shape the Cancer Genome.” Cell 155 (4): 948–62. https://doi.org/10.1016/J.CELL.2013.10.011.

Dewar, James M., and Johannes C. Walter. 2017. “Mechanisms of DNA Replication Termination.” Nature Reviews Molecular Cell Biology 18 (8): 507–16. https://doi.org/10.1038/nrm.2017.42.

Donley, Nathan, and Mathew J. Thayer. 2013. “DNA Replication Timing, Genome Stability and Cancer: Late and/or Delayed DNA Replication Timing Is Associated with Increased Genomic Instability.” Seminars in Cancer Biology 23 (2): 80–89. https://doi.org/10.1016/J.SEMCANCER.2013.01.001.

Elledge, S. J. 1996. “Cell Cycle Checkpoints: Preventing an Identity Crisis.” Science 274 (5293): 1664–72. https://doi.org/10.1126/science.274.5293.1664.

Filteau, Marie, Véronique Hamel, Marie-Christine Pouliot, Isabelle Gagnon-Arsenault, Alexandre K Dubé, and Christian R Landry. 2015. “Evolutionary Rescue by Compensatory Mutations Is Constrained by Genomic and Environmental Backgrounds.” Molecular Systems Biology 11 (10): 832. https://doi.org/10.15252/msb.20156444.

Fraser, Hunter B, Aaron E Hirsh, Lars M Steinmetz, Curt Scharfe, and Marcus W Feldman. 2002. “Evolutionary Rate in the Protein Interaction Network.” Science (New York, N.Y.) 296 (5568): 750–52. https://doi.org/10.1126/science.1068696.

Fumasoni, Marco, Katharina Zwicky, Fabio Vanoli, Massimo Lopes, and Dana Branzei. 2015. “Error-Free DNA Damage Tolerance and Sister Chromatid Proximity during DNA Replication Rely on the Polα/Primase/Ctf4 Complex.” Molecular Cell. https://doi.org/10.1016/j.molcel.2014.12.038.

Gaillard, Hélène, Tatiana García-Muse, and Andrés Aguilera. 2015. “Replication Stress and Cancer.” Nature Reviews Cancer 15 (5): 276–89. https://doi.org/10.1038/nrc3916.

Gambus, Agnieszka, Frederick van Deursen, Dimitrios Polychronopoulos, Magdalena Foltman, Richard C Jones, Ricky D Edmondson, Arturo Calzada, and Karim Labib. 2009. “A Key Role for Ctf4 in Coupling the MCM2-7 Helicase to DNA Polymerase Alpha within the Eukaryotic Replisome.” The EMBO Journal 28 (19): 2992–3004. https://doi.org/10.1038/emboj.2009.226.

Gisselsson, D, L Pettersson, M Höglund, M Heidenblad, L Gorunova, J Wiegant, F Mertens, P Dal Cin, F Mitelman, and N Mandahl. 2000. “Chromosomal Breakage-Fusion-Bridge Events Cause Genetic Intratumor Heterogeneity.” Proceedings of the National Academy of Sciences of the United States of America 97 (10): 5357–62. https://doi.org/10.1073/pnas.090013497.

Gourguechon, Stéphane, Liam J Holt, and W Zacheus Cande. 2013. “The Giardia Cell Cycle Progresses Independently of the Anaphase-Promoting Complex.” Journal of Cell Science 126 (Pt 10): 2246–55. https://doi.org/10.1242/jcs.121632.

Gresham, David, Michael M. Desai, Cheryl M. Tucker, Harry T. Jenq, Dave A. Pai, Alexandra Ward, Christopher G. DeSevo, David Botstein, and Maitreya J. Dunham. 2008. “The Repertoire and Dynamics of Evolutionary Adaptations to Controlled Nutrient-Limited Environments in Yeast.” Edited by Michael Snyder. PLoS Genetics 4 (12): e1000303. https://doi.org/10.1371/journal.pgen.1000303.

Hanna, Joseph S, Evgueny S Kroll, Victoria Lundblad, and Forrest a Spencer. 2001. “Saccharomyces Cerevisiae CTF18 and CTF4 Are Required for Sister Chromatid Cohesion.” Molecular and Cellular Biology 21 (9): 3144–58. https://doi.org/10.1128/MCB.21.9.3144-3158.2001.

Harcombe, WR, R Springman, and JJ Bull. 2009. “Compensatory Evolution for a Gene Deletion Is Not Limited to Its Immediate Functional Network.” BMC Evolutionary Biology 9 (1): 106. https://doi.org/10.1186/1471-2148-9-106.

Hollstein, M, D Sidransky, B Vogelstein, and C C Harris. 1991. “P53 Mutations in Human Cancers.” Science (New York, N.Y.) 253 (5015): 49–53. https://doi.org/10.1126/science.1905840.

Hughes, Timothy R., Christopher J. Roberts, Hongyue Dai, Allan R. Jones, Michael R. Meyer, David Slade, Julja Burchard, et al. 2000. “Widespread Aneuploidy Revealed by DNA Microarray Expression Profiling.” Nature Genetics 25 (3): 333–37. https://doi.org/10.1038/77116.

Jerison, Elizabeth R., and Michael M. Desai. 2015. “Genomic Investigations of Evolutionary Dynamics and Epistasis in Microbial Evolution Experiments.” Current Opinion in Genetics and Development 35: 33–39. https://doi.org/10.1016/j.gde.2015.08.008.

Koren, Amnon, Ilya Soifer, and Naama Barkai. 2010. “MRC1-Dependent Scaling of the Budding Yeast DNA Replication Timing Program.” Genome Research 20 (6): 781–90. https://doi.org/10.1101/gr.102764.109.

Koschwanez, John H., Kevin R. Foster, and Andrew W. Murray. 2013. “Improved Use of a Public Good Selects for the Evolution of Undifferentiated Multicellularity.” ELife 2 (January): e00367. https://doi.org/10.7554/eLife.00367.

Kouprina, N, E Kroll, V Bannikov, V Bliskovsky, R Gizatullin, A Kirillov, B Shestopalov, et al. 1992. “CTF4 (CHL15) Mutants Exhibit Defective DNA Metabolism in the Yeast Saccharomyces Cerevisiae.” Molecular and Cellular Biology 12 (12): 5736–47. http://www.ncbi.nlm.nih.gov/pubmed/1341195.

Laan, Liedewij, John H Koschwanez, and Andrew W Murray. 2015. “Evolutionary Adaptation after Crippling Cell Polarization Follows Reproducible Trajectories.” ELife 4 (October): e09638. https://doi.org/10.7554/eLife.09638.

Labib, Karim, and Agnieszka Gambus. 2007. “A Key Role for the GINS Complex at DNA Replication Forks.” Trends in Cell Biology 17 (6): 271–78. https://doi.org/10.1016/J.TCB.2007.04.002.

Lang, Gregory I., Daniel P. Rice, Mark J. Hickman, Erica Sodergren, George M. Weinstock, David Botstein, and Michael M. Desai. 2013. “Pervasive Genetic Hitchhiking and Clonal Interference in Forty Evolving Yeast Populations.” Nature 500 (7464): 571–74. https://doi.org/10.1038/nature12344.

Laughney, Ashley M., Sergi Elizalde, Giulio Genovese, and Samuel F. Bakhoum. 2015. “Dynamics of Tumor Heterogeneity Derived from Clonal Karyotypic Evolution.” Cell Reports 12 (5): 809–20. https://doi.org/10.1016/J.CELREP.2015.06.065.

Lengauer, Christoph, Kenneth W. Kinzler, and Bert Vogelstein. 1998. “Genetic Instabilities in Human Cancers.” Nature 396 (6712): 643–49. https://doi.org/10.1038/25292.

Levy, Sasha F., Jamie R. Blundell, Sandeep Venkataram, Dmitri A. Petrov, Daniel S. Fisher, and Gavin Sherlock. 2015. “Quantitative Evolutionary Dynamics Using High-Resolution Lineage Tracking.” Nature 519 (7542): 181–86. https://doi.org/10.1038/nature14279.

Li, Rong, and Andrew W. Murray. 1991. “Feedback Control of Mitosis in Budding Yeast.” Cell 66 (3): 519–31. https://doi.org/10.1016/0092-8674(81)90015-5.

Lind, Peter A, Andrew D Farr, and Paul B Rainey. 2015. “Experimental Evolution Reveals Hidden Diversity in Evolutionary Pathways.” ELife 4 (March). https://doi.org/10.7554/eLife.07074.

Liu, Gaowen, Mei Yun Jacy Yong, Marina Yurieva, Kandhadayar Gopalan Srinivasan, Jaron Liu, John Soon Yew Lim, Michael Poidinger, et al. 2015. “Gene Essentiality Is a Quantitative Property Linked to Cellular Evolvability.” Cell 163 (6): 1388–99. https://doi.org/10.1016/j.cell.2015.10.069.

Liu, Yuan, Thomas A Richards, and Stephen J Aves. 2009. “Ancient Diversification of Eukaryotic MCM DNA Replication Proteins.” BMC Evolutionary Biology 9 (1): 60. https://doi.org/10.1186/1471-2148-9-60.

Lynch, Michael, Way Sung, Krystalynne Morris, Nicole Coffey, Christian R Landry, Erik B Dopman, W Joseph Dickinson, et al. 2008. “A Genome-Wide View of the Spectrum of Spontaneous Mutations in Yeast.” Proceedings of the National Academy of Sciences of the United States of America 105 (27): 9272–77. https://doi.org/10.1073/pnas.0803466105.

Macheret, Morgane, and Thanos D Halazonetis. 2015. “DNA Replication Stress as a Hallmark of Cancer.” Annual Review of Pathology 10 (January): 425–48. https://doi.org/10.1146/annurev-pathol-012414-040424.

Mazouzi, Abdelghani, Alexey Stukalov, André C. Müller, Doris Chen, Marc Wiedner, Jana Prochazkova, Shih-Chieh Chiang, et al. 2016. “A Comprehensive Analysis of the Dynamic Response to Aphidicolin-Mediated Replication Stress Uncovers Targets for ATM and ATMIN.” Cell Reports 15 (4): 893–908. https://doi.org/10.1016/J.CELREP.2016.03.077.

McGeoch, Adam T, and Stephen D Bell. 2008. “Extra-Chromosomal Elements and the Evolution of Cellular DNA Replication Machineries.” Nature Reviews. Molecular Cell Biology 9 (7): 569–74. https://doi.org/10.1038/nrm2426.

Micco, Raffaella Di, Marzia Fumagalli, Angelo Cicalese, Sara Piccinin, Patrizia Gasparini, Chiara Luise, Catherine Schurra, et al. 2006. “Oncogene-Induced Senescence Is a DNA Damage Response Triggered by DNA Hyper-Replication.” Nature 444 (7119): 638–42. https://doi.org/10.1038/nature05327.

Michaelis, Christine, Rafal Ciosk, and Kim Nasmyth. 1997. “Cohesins: Chromosomal Proteins That Prevent Premature Separation of Sister Chromatids.” Cell 91 (1): 35–45. https://doi.org/10.1016/S0092-8674(01)80007-6.

Miles, J, and T Formosa. 1992. “Evidence That POB1, a Saccharomyces Cerevisiae Protein That Binds to DNA Polymerase Alpha, Acts in DNA Metabolism in Vivo.” Molecular and Cellular Biology 12 (12): 5724–35. http://www.pubmedcentral.nih.gov/articlerender.fcgi?artid=360512&tool=pmcentrez&rendertype=abstract.

Moyer, Stephen E, Peter W Lewis, and Michael R Botchan. 2006. “Isolation of the Cdc45/Mcm2-7/GINS (CMG) Complex, a Candidate for the Eukaryotic DNA Replication Fork Helicase.” Proceedings of the National Academy of Sciences of the United States of America 103 (27): 10236–41. https://doi.org/10.1073/pnas.0602400103.

Müller, Carolin A., Michelle Hawkins, Renata Retkute, Sunir Malla, Ray Wilson, Martin J. Blythe, Ryuichiro Nakato, et al. 2014. “The Dynamics of Genome Replication Using Deep Sequencing.” Nucleic Acids Research 42 (1): e3–e3. https://doi.org/10.1093/nar/gkt878.

Muñoz, Sergio, and Juan Méndez. 2016. “DNA Replication Stress: From Molecular Mechanisms to Human Disease.” Chromosoma, January, 1–15. https://doi.org/10.1007/s00412-016-0573-x.

Murray, A. 1994. “Cell Cycle Checkpoints.” Current Opinion in Cell Biology 6 (6): 872–76. https://doi.org/10.1016/0955-0674(94)90059-0.

Murray, a W. 1992. “Creative Blocks: Cell-Cycle Checkpoints and Feedback Controls.” Nature 359 (6396): 599–604. https://doi.org/10.1038/359599a0.

Neelsen, Kai J, Isabella M Y Zanini, Raquel Herrador, and Massimo Lopes. 2013. “Oncogenes Induce Genotoxic Stress by Mitotic Processing of Unusual Replication Intermediates.” The Journal of Cell Biology 200 (6): 699–708. https://doi.org/10.1083/jcb.201212058.

O’Donnell, Michael, Lance Langston, and Bruce Stillman. 2013. “Principles and Concepts of DNA Replication in Bacteria, Archaea, and Eukarya.” Cold Spring Harbor Perspectives in Biology 5 (7): a010108.. https://doi.org/10.1101/cshperspect.a010108.

Pai, Chen-Chun, and Stephen E Kearsey. 2017. “A Critical Balance: DNTPs and the Maintenance of Genome Stability.” Genes 8 (2). https://doi.org/10.3390/genes8020057.

Pardo, Benjamin, Laure Crabbé, and Philippe Pasero. 2017. “Signaling Pathways of Replication Stress in Yeast.” FEMS Yeast Research 17 (2): 1–11. https://doi.org/10.1093/femsyr/fow101.

Parker, Matthew W., Michael R. Botchan, and James M. Berger. 2017. “Mechanisms and Regulation of DNA Replication Initiation in Eukaryotes.” Critical Reviews in Biochemistry and Molecular Biology 52 (2): 107–44. https://doi.org/10.1080/10409238.2016.1274717.

Pasero, Philippe, and Alessandro Vindigni. 2017. “Nucleases Acting at Stalled Forks: How to Reboot the Replication Program with a Few Shortcuts.” Annual Review of Genetics 51 (1): 477–99. https://doi.org/10.1146/annurev-genet-120116-024745.

Payen, Celia, Sara C Di Rienzi, Giang T Ong, Jamie L Pogachar, Joseph C Sanchez, Anna B Sunshine, M K Raghuraman, Bonita J Brewer, and Maitreya J Dunham. 2014. “The Dynamics of Diverse Segmental Amplifications in Populations of Saccharomyces Cerevisiae Adapting to Strong Selection.” G3 (Bethesda, Md.) 4 (3): 399–409. https://doi.org/10.1534/g3.113.009365.

Poli, Jérôme, Olga Tsaponina, Laure Crabbé, Andrea Keszthelyi, Véronique Pantesco, Andrei Chabes, Armelle Lengronne, et al. 2012. “DNTP Pools Determine Fork Progression and Origin Usage under Replication Stress.” The EMBO Journal 31 (4): 883–94. https://doi.org/10.1038/emboj.2011.470.

Raghuraman, M. K., Elizabeth A. Winzeler, David Collingwood, Sonia Hunt, Lisa Wodicka, Andrew Conway, David J. Lockhart, Ronald W. Davis, Bonita J. Brewer, and Walton L. Fangman. 2001. “Replication Dynamics of the Yeast Genome.” Science 294 (5540): 115–21. https://doi.org/10.1126/SCIENCE.294.5540.115.

Rancati, Giulia, Jason Moffat, Athanasios Typas, and Norman Pavelka. 2018. “Emerging and Evolving Concepts in Gene Essentiality.” Nature Reviews Genetics 19 (1): 34–49. https://doi.org/10.1038/nrg.2017.74.

Rhodes, Jenny M, Miranda McEwan, and Julia A Horsfield. 2011. “Gene Regulation by Cohesin in Cancer: Is the Ring an Unexpected Party to Proliferation?” Molecular Cancer Research : MCR 9 (12): 1587–1607. https://doi.org/10.1158/1541-7786.MCR-11-0382.

Rojas Echenique, José I., Sergey Kryazhimskiy, Alex N. Nguyen Ba, and Michael M. Desai. 2019. “Modular Epistasis and the Compensatory Evolution of Gene Deletion Mutants.” Edited by Geraldine Butler. PLOS Genetics 15 (2): e1007958. https://doi.org/10.1371/journal.pgen.1007958.

Saayman, Xanita, Cristina Ramos-Pérez, and Grant W. Brown. 2018. “DNA Replication Profiling Using Deep Sequencing.” In Methods Mol Biol., 195–207. https://doi.org/10.1007/978-1-4939-7306-4_15.

Samora, Catarina P., Julie Saksouk, Panchali Goswami, Ben O. Wade, Martin R. Singleton, Paul A. Bates, Armelle Lengronne, et al. 2016. “Ctf4 Links DNA Replication with Sister Chromatid Cohesion Establishment by Recruiting the Chl1 Helicase to the Replisome.” Molecular Cell 0 (0): 121–34. https://doi.org/10.1016/j.molcel.2016.05.036.

Sarogni, Patrizia, Orazio Palumbo, Adele Servadio, Simonetta Astigiano, Barbara D’Alessio, Veronica Gatti, Dubravka Cukrov, et al. 2019. “Overexpression of the Cohesin-Core Subunit SMC1A Contributes to Colorectal Cancer Development.” Journal of Experimental & Clinical Cancer Research 38 (1): 108. https://doi.org/10.1186/s13046-019-1116-0.

Schultz, L. B., N. H. Chehab, A. Malikzay, Jr Di Tullio, E. S. Stavridi, and T. D. Halazonetis. 2000. “The DNA Damage Checkpoint and Human Cancer.” Cold Spring Harbor Symposia on Quantitative Biology 65: 489–98. https://doi.org/10.1101/sqb.2000.65.489.

Sharp, Nathaniel P, Linnea Sandell, Christopher G James, and Sarah P Otto. 2018. “The Genome-Wide Rate and Spectrum of Spontaneous Mutations Differ between Haploid and Diploid Yeast.” Proceedings of the National Academy of Sciences of the United States of America 115 (22): E5046–55. https://doi.org/10.1073/pnas.1801040115.

Siddiqui, Khalid, Kin Fan On, and John F X Diffley. 2013. “Regulating DNA Replication in Eukarya.” Cold Spring Harbor Perspectives in Biology 5 (9): a012930. https://doi.org/10.1101/cshperspect.a012930.

Simon, Aline C, Jin C Zhou, Rajika L Perera, Frederick van Deursen, Cecile Evrin, Marina E Ivanova, Mairi L Kilkenny, et al. 2014. “A Ctf4 Trimer Couples the CMG Helicase to DNA Polymerase Alpha in the Eukaryotic Replisome.” Nature 510 (7504): 293–97. https://doi.org/10.1038/nature13234.

Soulier, Jean, and Noel F. Lowndes. 1999. “The BRCT Domain of the S. Cerevisiae Checkpoint Protein Rad9 Mediates a Rad9–Rad9 Interaction after DNA Damage.” Current Biology. https://doi.org/10.1016/S0960-9822(99)80242-5.

Spencer, F., S. L. Gerring, C. Connelly, and P. Hieter. 1990. “Mitotic Chromosome Transmission Fidelity Mutants in Saccharomyces Cerevisiae.” Genetics 124: 237–49.

Steenwyk, Jacob L., Dana A. Opulente, Jacek Kominek, Xing-Xing Shen, Xiaofan Zhou, Abigail L. Labella, Noah P. Bradley, et al. 2019. “Extensive Loss of Cell-Cycle and DNA Repair Genes in an Ancient Lineage of Bipolar Budding Yeasts.” Edited by Sophien Kamoun. PLOS Biology 17 (5): e3000255. https://doi.org/10.1371/journal.pbio.3000255.

Szamecz, Béla, Gábor Boross, Dorottya Kalapis, Károly Kovács, Gergely Fekete, Zoltán Farkas, Viktória Lázár, et al. 2014. “The Genomic Landscape of Compensatory Evolution.” Edited by Nick H. Barton. PLoS Biology 12 (8): e1001935. https://doi.org/10.1371/journal.pbio.1001935.

Tanaka, H, Y Katou, M Yagura, K Saitoh, T Itoh, H Araki, M Bando, and K Shirahige. 2009. “Ctf4 Coordinates the Progression of Helicase and DNA Polymerase Alpha.” Genes to Cells : Devoted to Molecular & Cellular Mechanisms 14 (7): 807–20. https://doi.org/10.1111/j.1365-2443.2009.01310.x.

Tkach, Johnny M., Askar Yimit, Anna Y. Lee, Michael Riffle, Michael Costanzo, Daniel Jaschob, Jason A. Hendry, et al. 2012. “Dissecting DNA Damage Response Pathways by Analysing Protein Localization and Abundance Changes during DNA Replication Stress.” Nature Cell Biology 14 (9): 966–76. https://doi.org/10.1038/ncb2549.

Tsaponina, Olga, Emad Barsoum, Stefan U. Åström, and Andrei Chabes. 2011. “Ixr1 Is Required for the Expression of the Ribonucleotide Reductase Rnr1 and Maintenance of DNTP Pools.” PLoS Genetics 7 (5). https://doi.org/10.1371/journal.pgen.1002061.

Venkataram, Sandeep, Barbara Dunn, Yuping Li, Atish Agarwala, Jessica Chang, Emily R. Ebel, Kerry Geiler-Samerotte, et al. 2016. “Development of a Comprehensive Genotype-to-Fitness Map of Adaptation-Driving Mutations in Yeast.” Cell 166 (6): 1585-1596.e22. https://doi.org/10.1016/J.CELL.2016.08.002.

Villa, Fabrizio, Aline C. Simon, Maria Angeles Ortiz Bazan, Mairi L. Kilkenny, David Wirthensohn, Mel Wightman, Dijana Matak-Vinkovíc, et al. 2016. “Ctf4 Is a Hub in the Eukaryotic Replisome That Links Multiple CIP-Box Proteins to the CMG Helicase.” Molecular Cell 0 (0): 4601–5. https://doi.org/10.1016/j.molcel.2016.06.009.

Wang, H, D Liu, Y Wang, J Qin, and S J Elledge. 2001. “Pds1 Phosphorylation in Response to DNA Damage Is Essential for Its DNA Damage Checkpoint Function.” Genes & Development 15 (11): 1361–72. https://doi.org/10.1101/gad.893201.

Weinert, TA, and LH Hartwell. 1988. “The RAD9 Gene Controls the Cell Cycle Response to DNA Damage in Saccharomyces Cerevisiae.” Science 241 (4863): 317–22. https://doi.org/10.1126/science.3291120.

Wildenberg, Gregg A, and Andrew W Murray. 2014. “Evolving a 24-Hr Oscillator in Budding Yeast.” ELife 3 (January): e04875. https://doi.org/10.7554/eLife.04875.

Wilson, A C, S S Carlson, and T J White. 1977. “Biochemical Evolution.” Annual Review of Biochemistry 46 (1): 573–639. https://doi.org/10.1146/annurev.bi.46.070177.003041.

Xu, Huiling, Max Yan, Jennifer Patra, Rachael Natrajan, Yuqian Yan, Sigrid Swagemakers, Jonathan M Tomaszewski, et al. 2011. “Enhanced RAD21 Cohesin Expression Confers Poor Prognosis and Resistance to Chemotherapy in High Grade Luminal, Basal and HER2 Breast Cancers.” Breast Cancer Research 13 (1): R9. https://doi.org/10.1186/bcr2814.

Yao, Nina, and Mike O’Donnell. 2016. “Bacterial and Eukaryotic Replisome Machines.” JSM Biochemistry and Molecular Biology 3 (1): 1–7. http://www.ncbi.nlm.nih.gov/pubmed/28042596%0Ahttp://www.pubmedcentral.nih.gov/articlerender.fcgi?artid=PMC5199024.

Yona, Avihu H, Idan Frumkin, and Yitzhak Pilpel. 2015. “A Relay Race on the Evolutionary Adaptation Spectrum.” Cell 163 (3): 549–59. https://doi.org/10.1016/j.cell.2015.10.005.

Zegerman, Philip, and John F. X. Diffley. 2010. “Checkpoint-Dependent Inhibition of DNA Replication Initiation by Sld3 and Dbf4 Phosphorylation.” Nature 467 (7314): 474–78. https://doi.org/10.1038/nature09373.

Zegerman, Philip, and John F X Diffley. 2009. “DNA Replication as a Target of the DNA Damage Checkpoint.” DNA Repair 8 (9): 1077–88. https://doi.org/10.1016/j.dnarep.2009.04.023.

Zeman, Michelle K., and Karlene A. Cimprich. 2014. “Causes and Consequences of Replication Stress.” Nature Cell Biology 16 (1): 2–9. https://doi.org/10.1038/ncb2897.

Zheng, Dao-Qiong, Ke Zhang, Xue-Chang Wu, Piotr A Mieczkowski, and Thomas D Petes. 2016. “Global Analysis of Genomic Instability Caused by DNA Replication Stress in Saccharomyces Cerevisiae.” Proceedings of the National Academy of Sciences of the United States of America, November, 201618129. https://doi.org/10.1073/pnas.1618129113.

