## Supplementary figures for "The evolutionary plasticity of chromosome metabolism allows adaptation to DNA replication stress"

**Figure S1. Related to Fig. 1. (A)** Fitness of the *ctf4Δ* (yellow) and wt (grey) ancestors and of 16 evolved populations derived from them (8 each), relative to wt cells (*s*=0). Error bars represent standard deviations. Note that this panel shows the fitnesses of populations, whereas Fig. 1C shows the fitness of clones isolated from populations. **(B)** Bulk segregant analysis of evolved clones: One clone per population was crossed with a wt ancestor and subjected to bulk segregant analysis. For each clone, the mutations found to strongly segregate (>70%) with the evolved phenotype are reported. **(C)** Extract from Table S2: GO term that are enriched in the genes that were found to be either strongly significantly selected or segregating with the evolved phenotype by bulk segregant analysis.

**A**

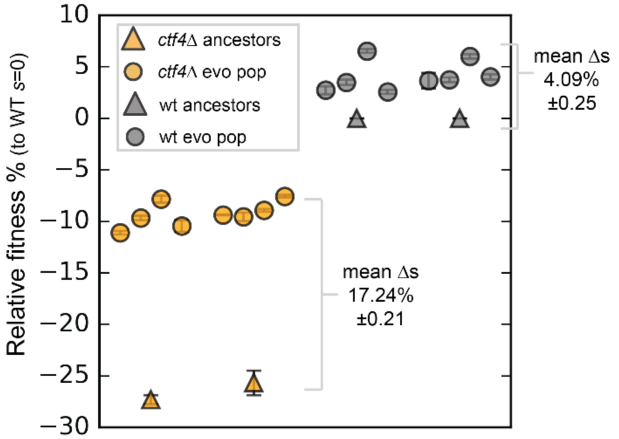


**B**

| Clone | Gene | Nucleotide change | AA and regulatory change | Segregation |
|---|---|---|---|---|
| EVO1-7 | *IXR1* | 1393 A→C | T465P | 97% |
| | *RAD9* | 3286 G→A | G1096E | 74% |
| | *TIR1* | 426-464del | 139-188del | 91% |
| EVO2-10 | *PSF3* | 134 G→A | S45N | 88% |
| | *NVJ2* | -559 G→C | promoter | 84% |
| EVO3-12 | *SIR4* | 1877 C→A | S626* | 97% |
| | *IXR1* | 922 C→T | Q308* | 94% |
| | *MMS1* | 2170 G→T | A724S | 88% |
| | *DPB11* | 1804 +A | S608* | 76% |
| | *RPS28B* | 42 -G | terminator | 90% |
| EVO4-2 | *IXR1* | 79 C→T | Q27* | 80% |
| | *SIR4* | 3140 C→T | S1047F | 95% |
| | *RAD9* | 2628 +A | K883* | 81% |
| | *RPS28B* | 42 -G | terminator | 76% |
| | *CDD1* | 68 +TTTT | terminator | 73% |
| EVO5-11 | *SLD5* | 388 G→A | E130K | 83% |
| | *CTH1* | 4 A→G | M2V | 71% |
| | *GIR2* | -197 T→C | promoter | 71% |
| EVO6-1 | *IXR1* | 1263 C→G | Y421* | 94% |
| | *SIR3* | 32 G→A | W11* | 79% |
| | *SMC2* | 940 C→A | R164S | 71% |
| | *UTR2* | -524 C→T | promoter | 80% |
| EVO7-7 | *PSF3* | 53 G→T | C18F | 93% |
| | *CTR9* | 976 T→A | L326I | 84% |
| | *DSF2* | 772 +A | T263* | 84% |
| EVO8-9 | *PSF1* | 599 T→A | I200N | 82% |

**C**

| #Term ID | Term description | Observed gene count | Background gene count | False discovery rate |
|---|---|---|---|---|
| GO:0006259 | DNA metabolic process | 13 | 499 | 0.00011 |
| GO:0006261 | DNA-dependent DNA replication | 7 | 117 | 0.00011 |
| GO:0006281 | DNA repair | 11 | 296 | 0.00011 |
| GO:0006302 | double-strand break repair | 7 | 131 | 0.00013 |
| GO:0051276 | chromosome organization | 11 | 566 | 0.0011 |
| GO:0007049 | cell cycle | 12 | 716 | 0.0019 |
| GO:0071103 | DNA conformation change | 5 | 117 | 0.0039 |
| GO:0006343 | establishment of chromatin silencing | 2 | 4 | 0.0041 |
| GO:0006310 | DNA recombination | 6 | 227 | 0.0073 |
| GO:0007076 | mitotic chromosome condensation | 2 | 11 | 0.0128 |
| GO:0006323 | DNA packaging | 3 | 56 | 0.02 |
| GO:0044773 | mitotic DNA damage checkpoint | 2 | 17 | 0.025 |
| GO:1903047 | mitotic cell cycle process | 6 | 310 | 0.0272 |

**A**

| Gene | Unique hits | Nucleotide change | AA change | Type | Note |
|---|---|---|---|---|---|
| *RAD9* | 5 | 2628 +A | frameshift | indel | K883* |
| *RAD9* | 1 | 3017 T→G | L1006W | substitution | |
| *RAD9* | 1 | 1904 +A | frameshift | indel | D638* |
| *RAD9* | 1 | 3287 G→A | G1096E | substitution | |
| *RAD9* | 1 | 3835 G→A | E1278K | substitution | |
| *MEC1* | 1 | 3917 C→T | A1306V | substitution | |
| *TEL1* | 1 | 2282 C→A | T2028K | substitution | kinase domain |
| *LCD1* | 1 | 536 G→A | R179H | substitution | |
| *DPB11* | 1 | 1804 +A | frameshift | indel | S608* |

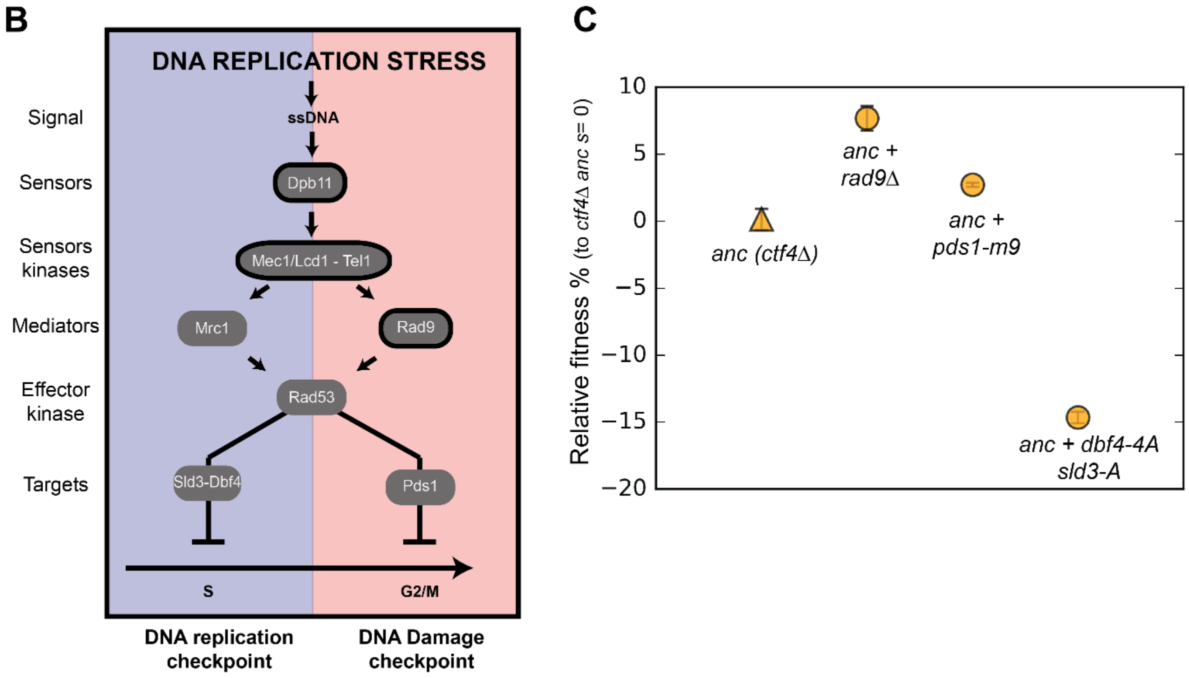


**Figure S2. Related to Fig. 2 (A)** Mutations affecting components of the DNA damage checkpoint that were found in evolved clones. **(B)** Schematic representation of the checkpoint signaling cascade induced by DNA replication stress. In blue (left) the DNA replication checkpoint, which delays S phase by phosphorylating Sld3 and Dbf4. In red (right) the DNA damage checkpoint, which induces a mitotic delay by phosphorylating Pds1. Factors highlighted in bold were found mutated in evolved lines. Some of the checkpoint factors (in the middle of the panel) play a role in both checkpoint responses, although this double role is likely not exerted simultaneously and may depend on the dynamics of the upstream checkpoint reactions. **(C)** The fitness of ancestral *ctf4*Δ strains carrying mutations affecting the DNA replication checkpoint signaling cascade at different levels, relative to the *ctf4Δ* ancestor (*s*=0). Error bars represent standard deviations.

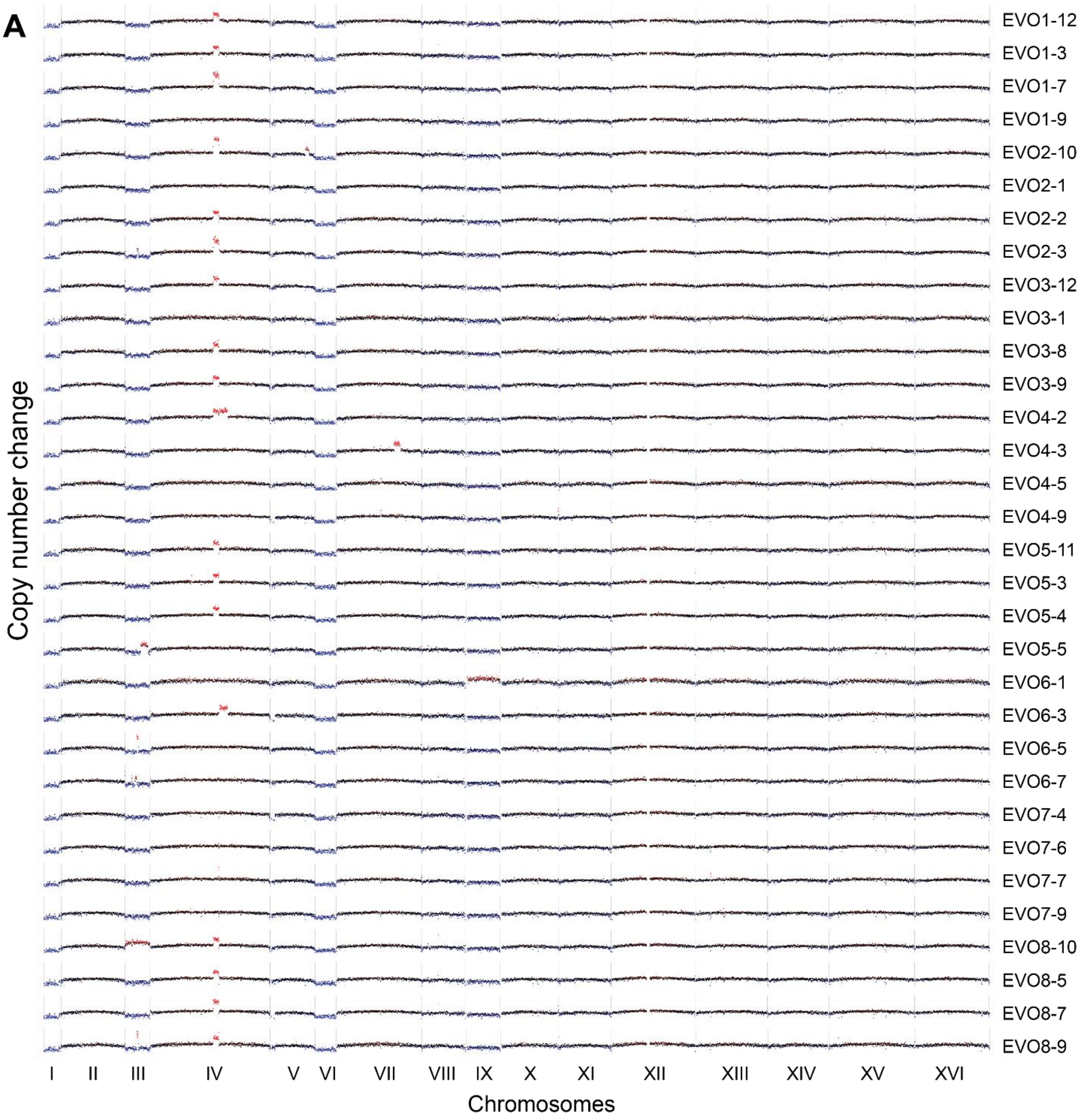
A
Copy number change
EVO1-12
EVO1-3
EVO1-7
EVO1-9
EVO2-10
EVO2-1
EVO2-2
EVO2-3
EVO3-12
EVO3-1
EVO3-8
EVO3-9
EVO4-2
EVO4-3
EVO4-5
EVO4-9
EVO5-11
EVO5-3
EVO5-4
EVO5-5
EVO6-1
EVO6-3
EVO6-5
EVO6-7
EVO7-4
EVO7-6
EVO7-7
EVO7-9
EVO8-10
EVO8-5
EVO8-7
EVO8-9
I II III IV V VI VII VIII IX X XI XII XIII XIV XV XVI
Chromosomes

48

**B**

| Gene | Unique hits | Nucleotide change | AA change | Type | Note |
|---|---|---|---|---|---|
| *RAD61* | 1 | 2628 T→A | Promoter | substitution | |
| *CHL1* | 1 | 2050 G→A | D684N | substitution | helicase domain |
| *PDS5* | 1 | 204 A→C | K68N | substitution | |
| *SMC2* | 1 | 940 C→A | R314S | substitution | |
| *TOF1* | 1 | 1244 C→A | P415Q | substitution | |
| *CSM3* | 1 | 370 G→C | V124L | substitution | |

**C**

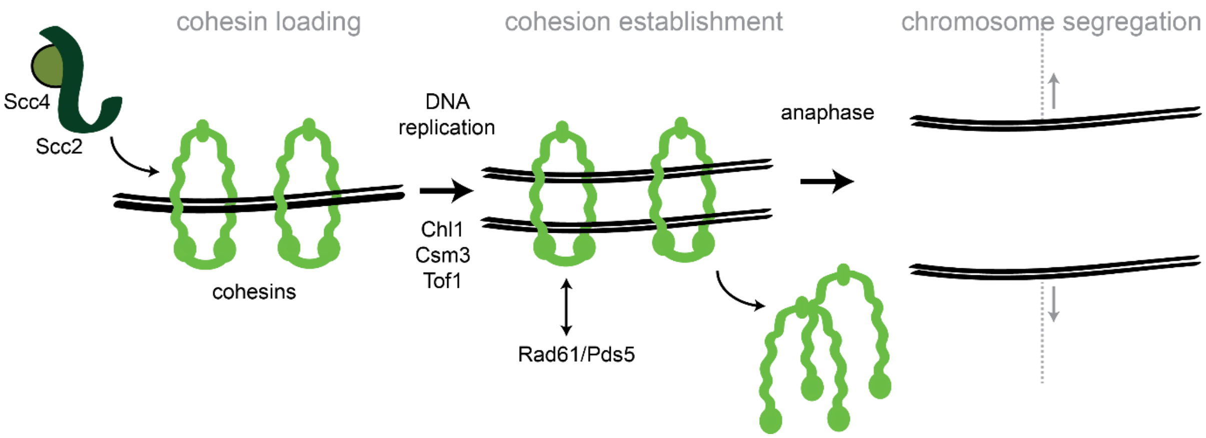


**Figure S3. Related to Fig. 3. (A)** CNVs affecting the genome of the 32 isolated evolved clones. Red highlights gains, blue highlights losses. Note that the aneuploidies affecting chromosome I, III, VI and IX, all of which are small chromosomes, may be due to the altered ancestral karyotype. We retrospectively found that ancestral *ctf4Δ* clones carried extra copies of these chromosomes, likely caused by chromosome mis-segregation acquired during strain construction or the initial pre-culture. Many evolved clones lose one of the two copies of these chromosomes during evolution arguing that aneuploidy for these chromosomes does not confer a long-term fitness advantage. **(B)** Mutations affecting genes implicated in chromosome segregation that were found in evolved clones. **(C)** Schematic representation of cohesion establishment: the cohesin loading complex (Scc2-Scc4) loads the cohesin ring onto chromosome in G1. With the passage of the replisomes during DNA replication, cohesion between sister chromatids is established. At the onset of anaphase, cohesin is cleaved to allow cells entering anaphase and segregating the chromosomes. Proteins whose genes were mutated in evolved strains are indicated next to the steps where they are believed to act.

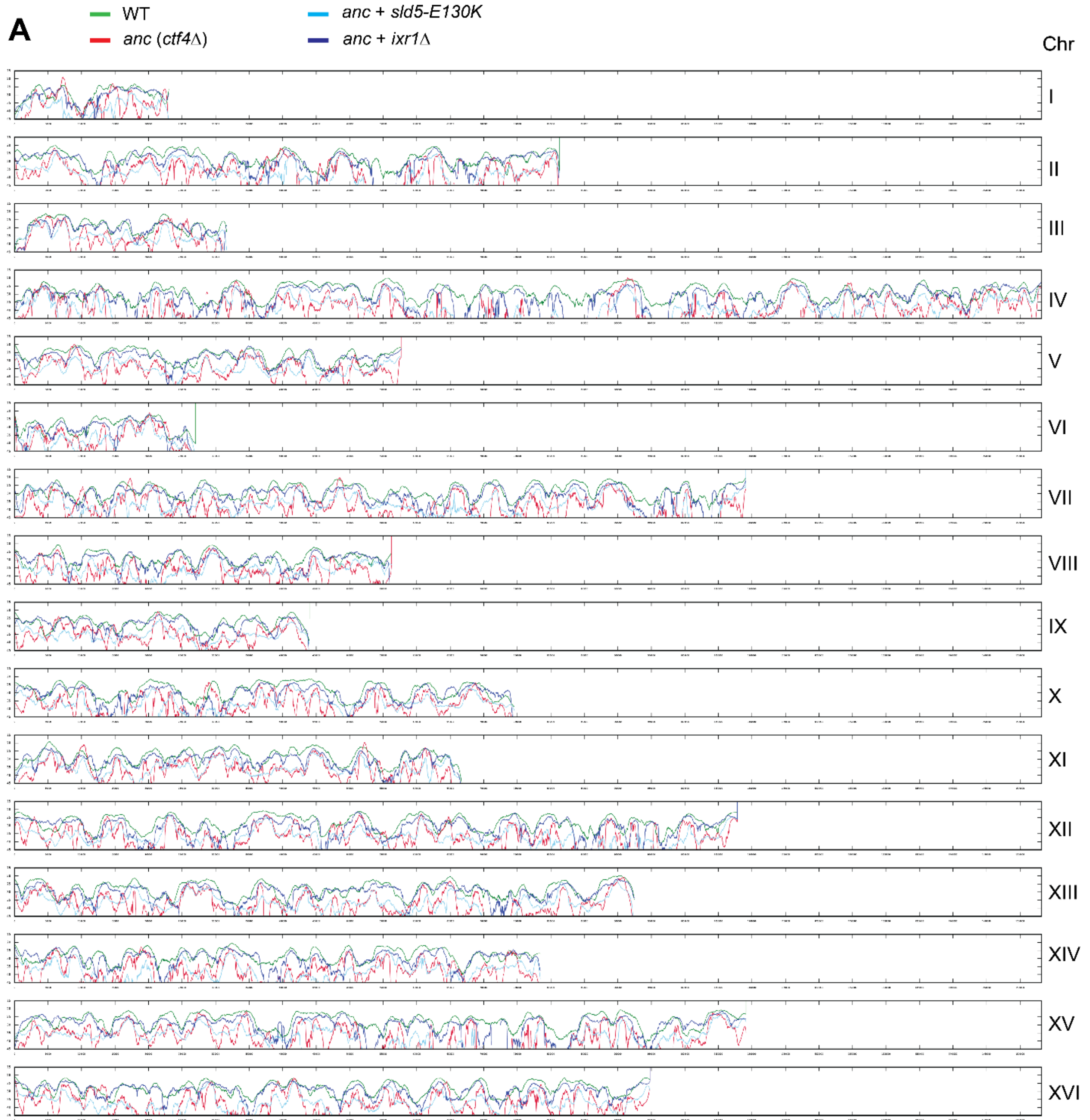
A
WT
anc (ctf4Δ)
anc + sld5-E130K
anc + ixr1Δ
Chr
I
II
III
IV
V
VI
VII
VIII
IX
X
XI
XII
XIII
XIV
XV
XVI

64

**B**

| Gene | Unique hits | Nucleotide change | AA change | Type | Note |
|---|---|---|---|---|---|
| *IXR1* | 1 | 1393 A→C | T465P | substitution | HMG domain |
| *IXR1* | 1 | 922 C→T | Q307* | substitution | |
| *IXR1* | 1 | 79 C→T | Q27* | substitution | |
| *IXR1* | 1 | 1263 C→G | Y421* | substitution | |
| *IXR1* | 1 | 1075 -C | Q359K | indel | R363* |
| *IXR1* | 1 | 994 C→T | Q332* | substitution | |
| *PSF3* | 1 | 569 G→A | W190* | substitution | |
| *PSF3* | 1 | 53 G→T | C18F | substitution | |
| *TOP1* | 1 | -243 G→A | promoter | substitution | |
| *TOP1* | 1 | 1257 A→T | L419F | substitution | catalytic core |
| *DPB2* | 1 | 64 T→C | Y22H | substitution | |
| *DPB2* | 1 | 1064 C→T | T355I | substitution | |
| *PSF1* | 1 | 599 T→A | I200N | substitution | |
| *DPB11* | 1 | 1804 +A | P602* | substitution | |
| *SLD5* | 1 | 388 G→A | E130K | substitution | |
| *CHL1* | 1 | 2050 G→A | D684N | substitution | helicase domain |
| *RFC1* | 1 | -156 +A | promoter | indel | |
| *TOF1* | 1 | 1244 C→A | P415Q | substitution | |
| *CSM3* | 1 | 370 G→C | V124L | substitution | |

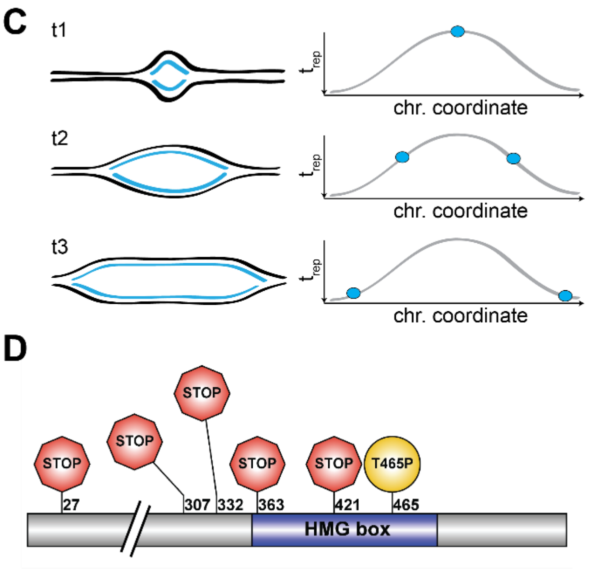


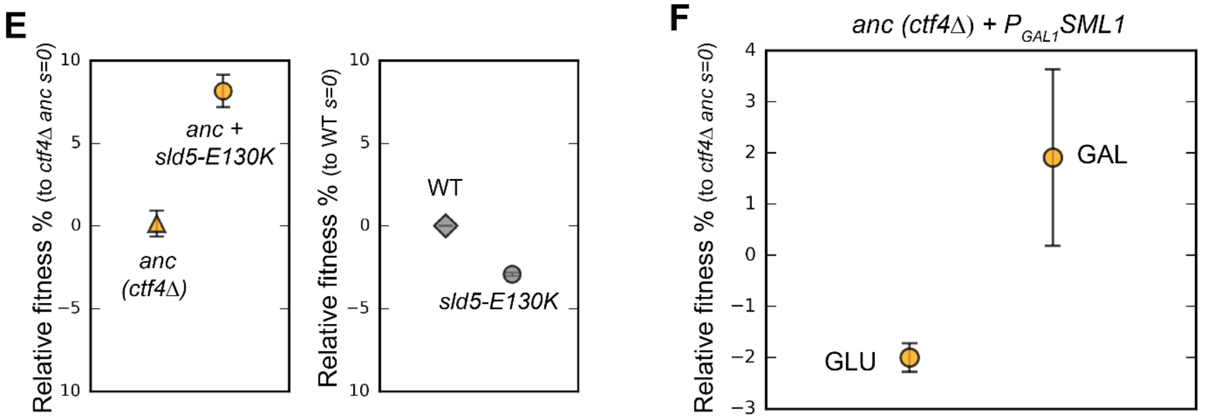


**Figure S4. Related to Fig. 4. (A)** Genome-wide DNA replication profiles of wt, the *ctf4*Δ ancestor, and two double mutant strains: *ctf4Δ sld5-E130K* and *ctf4Δ ixr1Δ*. $t_{rep}$ refers to the time at which 50% of the cells in the population replicated a region. $t_{rep}$ was derived from the change in DNA copy numbers over time, measured by deep sequencing (see material and methods). **(B)** Mutations affecting genes implicated in DNA replication that were found in evolved clones. **(C)** Schematic representation of the two replication forks arising from an origin of replication, and the related signal they generate in the replication profiles. **(D)** Mutations affecting Ixr1 found in evolved clones. Note that one stop codon (Q332*) resulted from an upstream frameshift. **(E)** Fitness effect of *sld5-E130K* on *ctf4Δ* ancestor cells (left panel) and on wt, *CTF4* cells (right panel). The fitness measurements are relative to *ctf4Δ* and wt respectively. Error bars represent standard deviations. **(F)** Effect of altered levels on deoxyribonucleotide triphosphates (dNTPs) on ancestor cells. Error bars represent standard deviations. *ctf4Δ* ancestor cells carrying a conditional $P_{GAL1}$-*SML1* allele were used. Sml1 is an inhibitor of the ribonucleotide reductase, an enzyme essential for dNTP production. *SML1* was expressed from the *GAL1* promoter, that is inhibited by glucose and strongly activated by galactose. A *ctf4Δ* $P_{GAL1}$-*SML1* strain was pre-cultured in YP + 2% raffinose and then competed against a *ctf4Δ* reference strain either in YP + 2% glucose (left side), or in YP + 2% galactose 2% raffinose (right side). This should result in dNTP overproduction (glucose) and shortage (galactose).

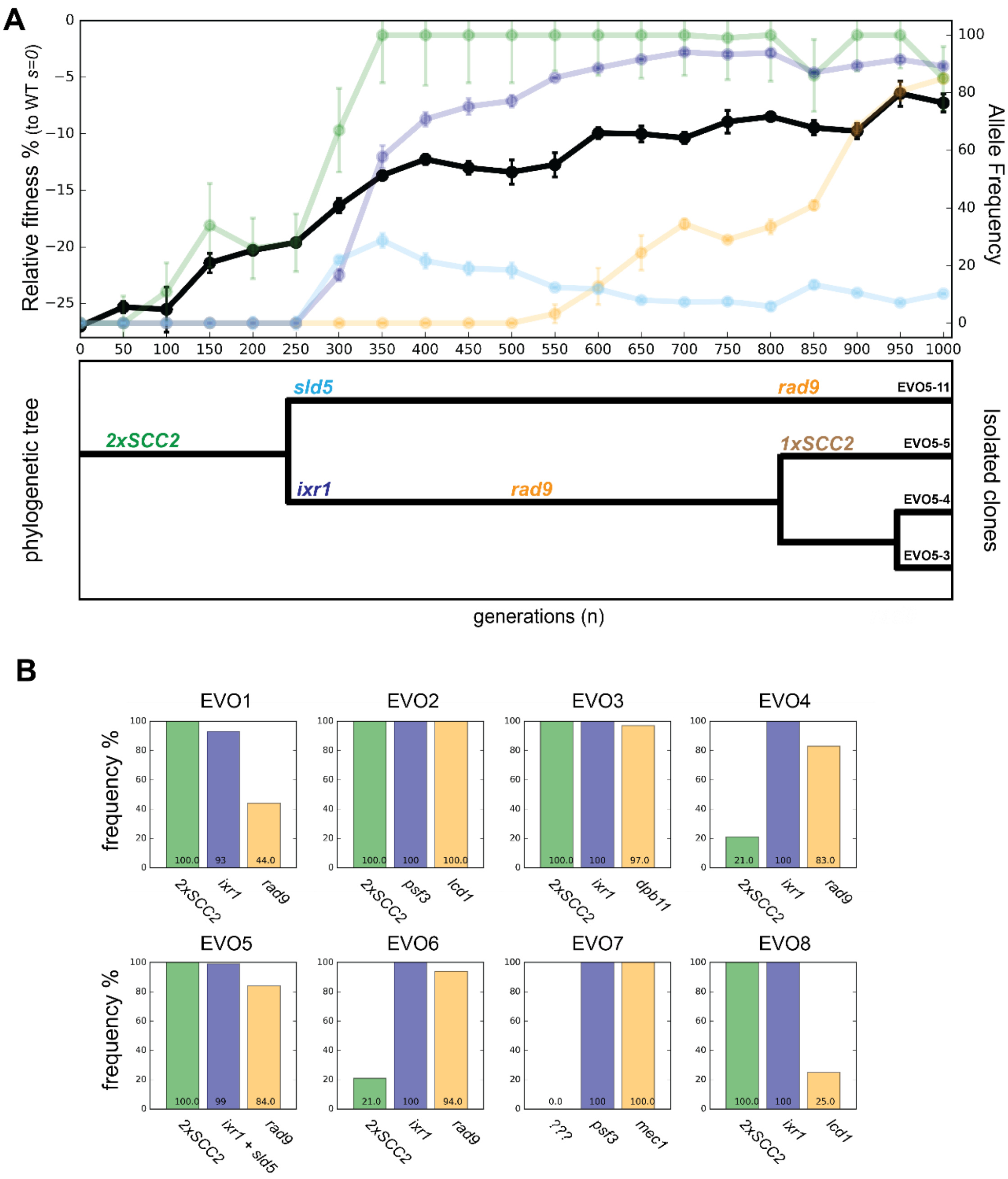


**Figure S5. Related to Fig. 5. (A)** Fitness of population EVO5 relative to wt (*s*=0) measured every 50 generations during the experiment (upper panel, dark plot, left y axis) with the frequency of mutant alleles included for reference (upper panel, faint plots, right x axis). Error bars represent standard deviations. Phylogenetic tree for clones isolated from population 5 (lower panel). Linkage was derived from analyzing whole genome sequences of the individual clones (TableS1), while branch length was inferred from the allele frequencies obtained by Sanger sequencing. **(B)** Frequencies of putative adaptive mutations in the cohesion, replication and checkpoint modules in the evolved populations at the conclusion of the experiment (generation 1000). The putative adaptive mutations were inferred based on results obtained for population EVO5. When the experimentally validated genes were not present, closely interacting genes were considered. Alleles frequencies in populations were obtained by deep sequencing of genomic DNA extracted from a population sample.
